# Large-scale proteogenomics characterization of the *Mycobacterium tuberculosis* hidden microproteome

**DOI:** 10.1101/2023.11.26.568715

**Authors:** Eduardo V. de Souza, Pedro F. Dalberto, Adriana C. Miranda, Alan Saghatelian, Antonio Michel Pinto, Luiz A. Basso, Pablo Machado, Cristiano V. Bizarro

## Abstract

Tuberculosis remains a burden to this day, due to the rise of multi and extensively drug-resistant bacterial strains. The genome of *Mycobacterium tuberculosis (Mtb)* underwent an annotation process that excluded small Open Reading Frames (smORFs), which encode a class of peptides and small proteins collectively known as microproteins. As a result, there is an overlooked part of its proteome that is a rich source of potentially essential, druggable molecular targets. Here, we employed our recently developed proteogenomics pipeline to identify novel microproteins encoded by smORFs in the genome of *Mtb* usings hundreds of mass spectrometry experiments in a large-scale approach. We found protein evidence for hundreds of novel microproteins and identified smORFs potentially involved in bacterial growth and virulence. Moreover, many smORFs are co-expressed or share operons with a myriad of biologically relevant genes and may play a role in antibiotic response. Together, our data presents a resource of unknown genes that play a role in the success of *Mtb* as a widespread pathogen.

## Introduction

A large portion of the bacterial proteome has been overlooked for as long as pipelines try to annotate genomes. One of the usual limitations of pure genomics approaches is the lack of confidence to accurately predict that an Open Reading Frame (ORF) shorter than 300 codons (smORF) is indeed encoding a microprotein, and does not represent one of the many artifacts following the inclusion of hundreds of thousands putative sequences into the analysis, which led bioinformaticians to establish an arbitrary cutoff of 300 codons to improve the accuracy of genome annotation ^1,2^. Furthermore, one of the reasons that may hamper smORF identification when using homology-based approaches for gene annotation is the fact that many smORFs are actually the outcome of a *de novo* gene birth event, which makes it unlikely for such strategy to consider them for the subsequent annotation steps ^3–6^. The field of proteogenomics, which employs different omics techniques alongside genomics, such as transcriptomics and proteomics, has been useful for a wide array of analyses, and can also be used to support genome annotation ^7^. Using Mass Spectrometry (MS) data to search a database containing the whole coding potential of a genome or transcriptome, different studies have reported the identification of novel bacterial microproteins encoded by smORFs using this unbiased approach ^8,9^.

One of the problems that surround proteogenomics analyses is their low sensitivity when compared to other methods like Ribosome Profiling, which is able to identify thousands of translated smORFs in some cases ^10–12^. There is a concern about how noisy such data might be, though, as the overlap of the results coming from different pipelines for downstream analysis seems to be quite small ^13^. Using proteomics, not only we get evidence at the protein instead of the translational level, but the result is also usually more reliable than using the ribosome profiling evidence alone, as there are findings showing pervasive translation all across the transcriptome ^10^, which could mean that, even though translation is occurring, there is no way of telling whether the protein originating from this translation event is stable or not without further experimentation. To overcome the limitations of proteomics regarding sensitivity, we gathered 680 MS experiments from different studies from our group, and used them all to search for possible smORF-encoded peptides (SEPs) that might be hidden in the proteome of *Mycobacterium tuberculosis* (*Mtb*), the etiological agent of tuberculosis. This is a straightforward but powerful approach to increase the chance of identifying novel, but rather inconspicuous sequences, simply by increasing the number of samples that might contain these microproteins. In the case of bacteria, using proteogenomics also seems to have better potential to yield good results than it does for eukaryotic genomes. One of the reasons is the much smaller genome, which results in considerably smaller databases, reducing the number of false negatives by decreasing the stringency when assessing the False Discovery Rate (FDR).

Using our recently published pipeline, µProteInS ^14^, we were able to uncover a vast amount of novel microproteins encoded by the genome of *Mtb*. We also characterized microproteins sharing different degrees of confidence, and extracted meaningful biological information from the newly-uncovered sequences, such as conserved protein domains, expression and co-expression levels across different conditions. We also demonstrate the occurrence of translation of smORFs delimited by the same stop but different start codons. Using this open-architecture approach, we have also found evidence for smORFs coming from *de novo* gene birth events in Mycobacterial evolution. Moreover, our exploratory analysis revealed that some of these smORFs seem to be embedded by genomic segments that either affect growth or are essential for bacterial survival. Thus, these novel coding sequences represent interesting candidates for future and more in-depth experimental studies that focus on the biological roles of mycobacterial microproteins.

## Results

μProteInS identifies novel microproteins encoded by smORFs using both the genome and the transcriptome as proteomics databases. During the first step of the pipeline, *assembly*, we assembled the transcriptome using a reference-guided approach, which borrows aspects of a *de novo* transcriptome assembly, allowing the identification of both known and novel transcripts. This was done using the raw strand-specific RNA-Seq reads for the control samples from the study E-MTAB-1616. With the three frame translation of the transcript sequences using four different start codons (ATG, ATT, GTG, and TTG), we generated a custom database from this assembly containing 76353 putative microproteins. The database resulting from the six frame translation of the reference genome contained 217657 microprotein sequences. 5609 sequences inside both databases consist of already annotated proteins from UniProt, NCBI RefSeq and Mycobrowser. Both of these databases were used for the *ms* step of µProteInS, which performs the peptide search against MS data. We used 680 MS experiments to search spectral data against two custom databases, generated from the six and three-frame translations of the reference genome and the assembled transcriptome, respectively. These results, after undergoing *postms*, the post-processing step of µProteInS, yielded more than 959 microproteins in total for the genome database, at a FDR of 0.01. This seems to be egregiously high, as the genome of *Mtb* currently encodes 4173 genes, as stated in Mycobrowser. Using the last mode of µProteInS, *validate*, which employs a random forest model pre-trained on manually inspected data, we classified the Peptide-Spectrum Matches (PSMs) as either High (HC) or Low Confidence (LC). After this validation step, we were able to identify 46 smORFs from the genome database (gORFs) and 26 smORFs from the transcriptome database (tORFs) with at least one HC spectrum matching their encoded microproteins (HC smORFs). These microproteins are our most reliable identifications, and henceforth will be treated as our gold standard, but this does not mean that the remaining portion of the results does not contain reliable identifications. Although HC smORFs possess higher quality MS spectra, many smORFs appeared in a considerable amount of replicates, which led us to consider them for the subsequent analyses as well. Out of the annotated proteins from the concatenated proteome from three databases, we could identify 4676 from the genome (83.3%), and 4364 from the transcriptome (77.8%) databases, to a total of 4762 non-redundant proteins from both databases (84.8%).

**Figure 1.**
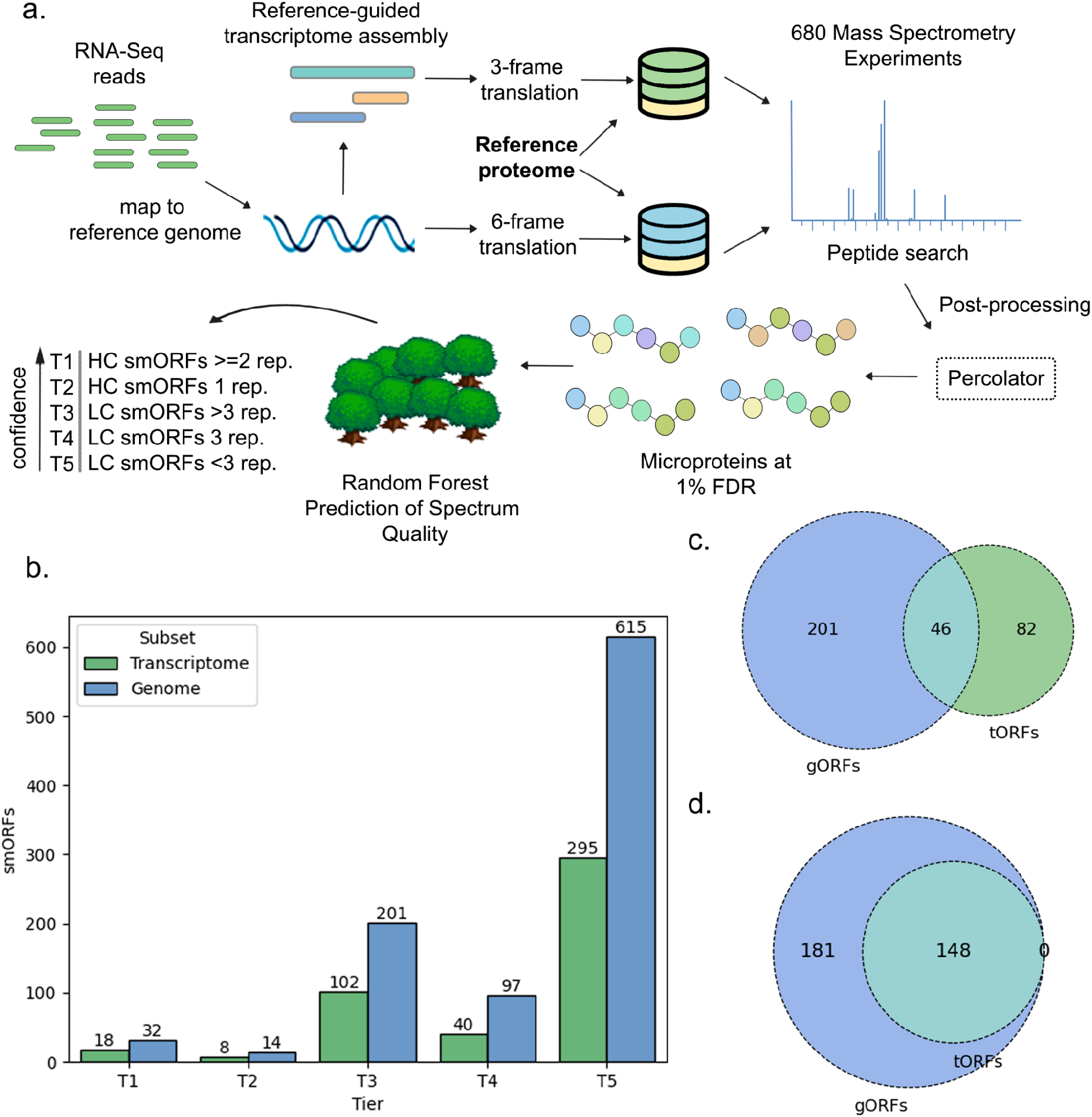
Proteogenomics identification of novel microproteins. **a,** Proteogenomics workflow to identify and characterize novel microproteins encoded by smORFs in *Mtb.* The workflow incorporates µProteInS pipeline into a large-scale approach to characterize Mass Spectrometry evidence by different degrees of confidence. Following the standard μProteInS pipeline, RNA-Seq reads are used to assemble the transcriptome, then custom proteomics databases from both the genome and transcriptome are translated *in silico* to search for the whole coding potential of *Mtb*. This is done in the third mode of μProteInS, where a large number of experiments were searched and scored. The last mode validates the findings, predicting High-Confidence (HC) and low-confidence (LC) spectra. The findings are then grouped and ranked into tiers regarding their level of confidence across multiple replicates, where lower tiers indicate smORFs with stronger evidence. **b,** Number of novel smORFs identified for each Tier coming from both the genome (gORFs) and transcriptome (tORFs) databases. **c,** Intersection between the results of gORFs and tORFs up to T3. **d,** Intersection between gORFs and tORFs after classifying as an intersection any smORF that had evidence for one group, but was absent for the other while being present in the initial database due to an exclusion because of the bigger Decoy database.

To classify these smORFs by different degrees of confidence, by making use of our extensive dataset, we divided them into a hierarchical classification of Tiers, where lower Tier numbers represent identifications with higher reliability (figure 1A). There are five Tiers: HC smORFs that appeared in two or more replicates (T1), HC smORFs appearing in only one replicate (T2), smORFs that have no HC PSM (LC smORFs), but appear in more than three replicates (T3), LC smORFs that appear in exactly three replicates (T4), and LC smORFs that appear in less than three replicates (T5). With this classification, we found 32 T1 gORFs and 18 T1 tORFs, and 14 T2 gORFs and 8 T2 tORFs (fig. 1B). This is our gold standard, but we decided to consider for the subsequent analyses all the smORFs up to T3, since the ones in this group appear in many replicates, passed the traditional filter of 1 % FDR, and could still represent important findings. T3 comprises the highest number of novel smORFs, while this number decreases for T4, which sits at 97 gORFS and 40 tORFs, and goes up to enormous amounts at T5. The latter most likely comprises lots of artifacts that do not account for real sequences, but it may still be informative when comparing different characteristics among these groups.

### Using a transcriptome alongside the genome increases the number of identifications

Given the lack of alternative splicing on the bacterial transcriptome, the common approaches for bacterial proteogenomics usually rely solely on the six frame translation of the genome for the generation of custom databases for proteomics ^8,9,15^. This would not be ideal for eukaryotes, due to bloated databases resulting from an increased genome size, alternative splicing events, and many long, repetitive intergenic regions carrying multiple regulatory elements. In bacteria, however, the six-frame translation approach does not face these limitations. There are still many advantages for including a transcriptome assembly into the workflow, nevertheless. As shown in figure 1C, many microproteins could only be identified using the transcriptome database. This is not a result of a transcript isoform with an exon organization that could not be retrieved during the six frame translation of the genome - rather, we hypothesized that this happened due to the fact that the transcriptome database is almost three times smaller than its genomic counterpart, which results in a less stringent q-value cutoff when assessing the FDR. This largely increases the number of identifications: up to T3, the transcriptome database search yielded 82 unique tORFs, which would have been missed if we used the genome alone for the *database* step of µProteInS. To test this, we checked which ones of these unique tORFs were present in the genome database originally, but were discarded during FDR assessment. For all sets of tiers (fig. 1D), when including as an intersection any identification that was initially present in both databases, we found out that all these unique tORFs were indeed an outcome of the usage of a smaller database, as we used the same Mass Spectrometry datasets to perform the peptide search for both databases. This could mean that, although databases generated from prokaryotic databases are less affected by bloated Mass Spectrometry databases than eukaryotes, they can still benefit from database shrinking strategies, like the generation of a custom database from the three-frame translation of the transcriptome.

**Figure 2.**
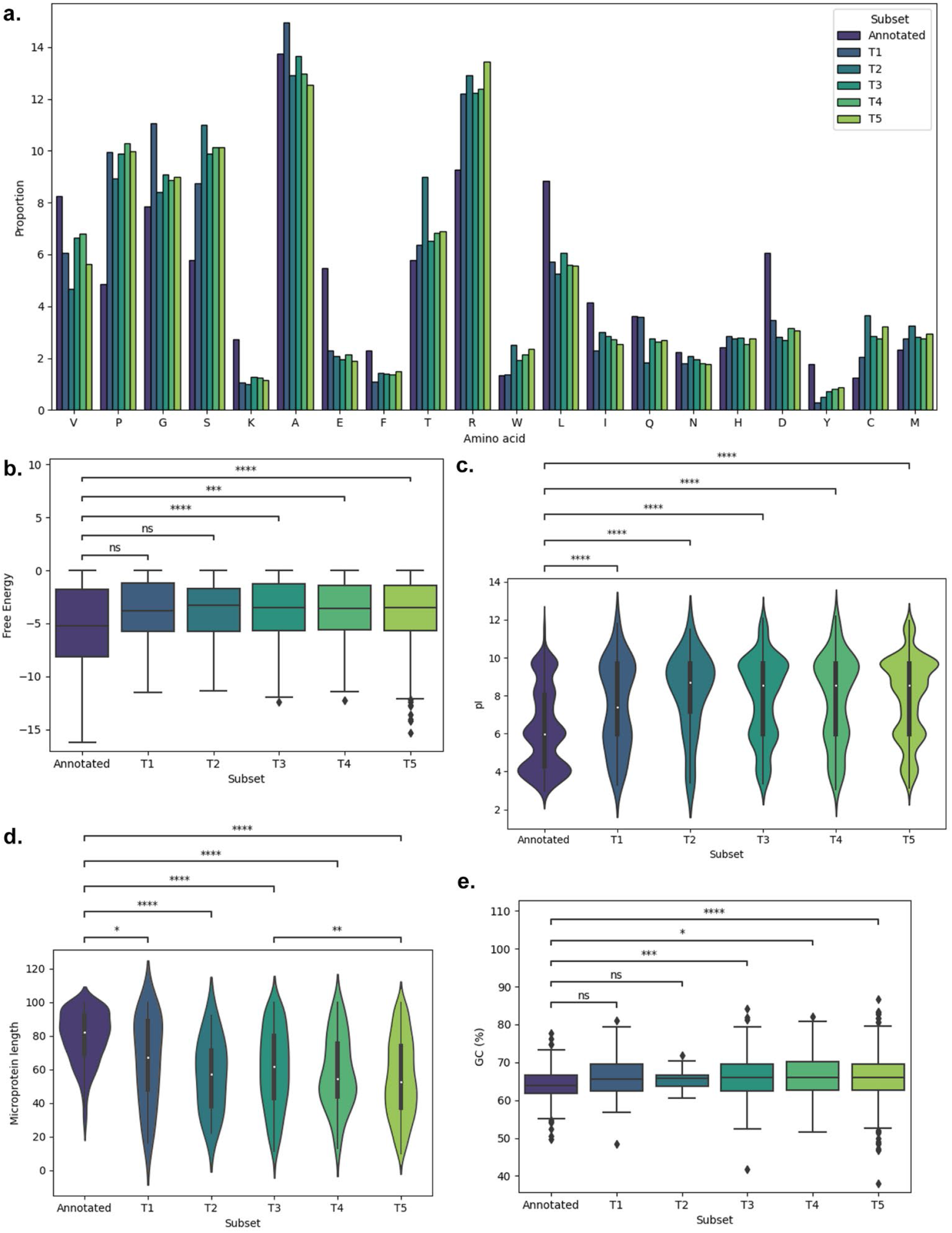
Characterization of smORFs and their encoded microprotein sequences across multiple tiers. **a,** Amino acid distribution of the novel smORFs and annotated genes. **b,** Free energy levels of the upstream sequences of the novel smORFs and annotated genes when binding to the 16S rRNA of H37Rv. **c,** Distribution of isoelectric points of the tryptic peptides coming from digested novel and annotated microproteins. **d,** Overall novel and annotated microprotein length. **e,** GC percentage of the nucleotide sequences of novel and annotated smORFs.

The inclusion of a transcriptome assembly, be it *de novo* or reference-guided, also brings other advantages to a proteogenomics workflow. The most important one is the addition of another layer of evidence to the finding, as tORFs have three different layers of evidence, at the gene, transcript, and protein level. When translating the nucleotide sequences from expressed transcripts, there is less room for artifacts coming from intergenic regions. Also, the read coverage of a RNA-Seq experiment for a smORF is usually much higher than the number spectral counts for its microprotein, which makes it easier to quantify these sequences and opens up the possibility to perform analysis like differential gene expression at the transcript level.

### smORF and microprotein sequence characterization

Seeking to understand where these smORFs differ from previously known ORFs, as this could pinpoint the characteristics that might be precluding their identification by traditional methods, we analyzed the GC content of the smORFs and the amino acid distribution of the novel microproteins, since they could present some sequence bias, as studies have shown before ^5,11^. Indeed, many amino acids differ in their abundance among the novel sequences when compared to the annotated proteins in our reference proteome database. A few amino acids have a clear prevalence in the sequence of annotated proteins, such as valine, lysine, glutamate, leucin, aspartate, and tyrosine (Fig. 2A). Lysine is of utmost importance for peptide identification by Mass Spectrometry-based proteomics, since trypsin, the most widely used enzyme for protein digestion, uses this amino acid as one of its cleavage sites ^16^. Thus, fewer lysines result in fewer tryptic peptides, which could hamper the identification of these microproteins. Some amino acids are more common in the novel sequences, though, such as proline, arginine, and cysteine. This sequence bias might be one of the reasons behind the poor performance of traditional algorithms for inferring conserved protein domains from microproteins, as those models were all trained based on the current literature at the time, which mostly disregarded these short sequences. Consequently, a not-so-accurate prediction would be expected when trying to generalize from these models. If we look at the GC content of their coding sequences, the novel smORFs from T1 and T2 present no significant statistical difference from the annotated CDSs, but smORFs from T3 to T5 do, with the biggest difference shared by T5 smORFs, which have the lowest level of confidence among the novel identifications. This sequence bias also affects the isoelectric point of the protein (fig 2c), one of the key physicochemical properties of a peptide for MS-based proteomics, where peptides coming from the digestion of novel microproteins tend to have a higher isoelectric point. The microprotein length varies drastically when compared to the median length of the annotated proteins, as novel microproteins from all tiers have a very strong tendency for shorter sequences (p < 0.001). As a result, the universe of possible tryptic peptides coming from each protein is also smaller, which hampers their identification by MS-based proteomics.

Ribosome-binding sites (RBS) are necessary for translation initiation in many mRNAs, and in bacteria they may appear multiple times in the same polycistronic transcript, located upstream of different ORFs. We sought to check which of the smORF-containing transcripts (SCTs) carried a RBS, as this would increase their chance of being translated. To do so, we extracted the 21 nucleotides upstream from each smORF in their respective SCTs and checked the free energy level resulting from the binding of this sequence to the 3’ end of the 16S rRNA of *Mtb.* In general, RBS of annotated ORFs have lower free energy levels than most of the novel smORFs. In fact, when using the thresholds defined by Starmer et al (2006) ^17^, we found out that only a small portion of the novel smORFs carry an upstream sequence that could be classified as a RBS. This is not surprising, as ORFs lacking a Shine-Dalgarno sequence are very common in the transcriptome of Mycobacteria. Moreover, even in eukaryotes, the Kozak sequence seems to be absent for many smORFS in SCTs ^18^.

### Conserved protein domains are identified within the sequences of novel microproteins

We Identified a variety of protein domains with different functions searching all the sequences with InterPro. The vast majority of the novel microproteins have at least one intrinsic disorder domain, which is common for all gORFs and tORFs from T2. 80.9% and 73.08% of all domains in gORFs and tORFs from Tier 1, respectively, have intrinsic disorder, whilst 96.21%, 98.53% and 96.26% of domains in gORFs from T3 onwards and more than 98% of domains in tORFs from T3 and forth have these same domains. We found several other domains such as ribosomal proteins in 14.28% of gORFs and 23.1% of tORFs T1, as well as prokaryotic membrane lipoprotein lipid attachment site profiles, which we found in 3.85% of T1 smORFs’ domains. In addition, we found a tRNA translation domain in one of the gORFs T3 microproteins, and we finally identified the biggest divergence of domains in T5, including YggU-like, opposite-ABC transporters and antisense-to-16S rRNA domains. Summing up, we found that most of the microproteins have more than one domain, mainly those in gORFs T1, with a total of 42 domains in 21 microproteins. We found the same proportion in gORFs T5, with 454 domains in 293 sequences. To identify the presence of any transmembrane helices within the new microproteins, we searched sequences from Tier 1 up to Tier 3 using both Phobius and TMHMM as predictors. 11 microproteins from T1 and T3 had a prediction of transmembrane helices above 50% with Phobius and gORFs 18752, 26725 and 65479 had transmembrane predictions on both Phobius and TMHMM. It is also worth noting that we found 15 smORFs with internal helices identified by Psortb as well as transmembrane predictions by TMHMM or Phobius. We found gORF 68556 was the only smORF with signal peptides characterized by SignalP but we also identified disorder, transmembrane and lipid attachment site domains. Complete data for domain prediction is available in Supplementary Table 1. When we ran CELLO on all smORFs up to tier 3, we only found gORF_95002 and tORF_gene-Rv2975a with GO terms, and both had ribosomes as cellular components. Moreover, we identified 271 smORFs from Tier 1 to Tier 3 with GO terms using DeepGoWeb. We encountered 13 different cellular components, with almost 50% of them being cellular anatomical entities; 17 different biological processes, out of which 42% were biological regulation, followed by cellular process and regulation of cellular process; and binding and catalytic activity as molecular functions.

### Genomic and transcriptomic landscape of the mycobacterial smORFome

After the identification of the novel microproteins encoded by smORFs, we wanted to understand how they are distributed and interconnected across the mycobacterial genome. To illustrate this, we generated a circos plot showing the locations of the novel smORFs in a genome-wide level (figure 3a). The plot tells us that the smORFs are not clustered in a specific region, but widespread all across the genome of the bacterium, sometimes filling gaps in its few intergenic regions. The third ring also shows the overall level of conservation of these smORFs in closely-related bacteria, suggesting strong evidence for *de novo* gene birth events. The links in the center of the plot also show how important and interconnected these novel sequences are for the transcriptome, as they represent evidence of co-expression between one smORF and an annotated gene, suggesting strong co-regulation of smORFs and canonical genes.

### Novel smORFs overlap the genomic loci of annotated genes and essential regions for bacterial survival

Overlooked outside viruses until recently, overlapping genes have growing evidence to be more common than initially thought, as 27% of prokaryotes coding sequences (CDS) seem to overlap another CDS ^19^. Indeed, almost all of the novel smORFs we found overlap an annotated region in the genome (Supplementary Table 2), while only 1.24% are found within intergenic regions. We defined any smORF within 500 bp up or downstream from an annotated gene as an upstream ORF (uORF) or downstream ORF (dORF), respectively. This is important as we are predicting sequences from the genome as well, so it is still possible to classify an overlap for a smORF that could not be found within the transcriptome assembly. The rarity of smORFs within intergenic regions is probably due to the compact genome of *Mtb*, where most genes are very close to each other. The majority of overlapping cases comprise smORFs that are nested out of frame, nested in the opposite strand, or overlapping an annotated gene (Fig 2a). Although most of the features overlapped by a smORF are CDSs, we could also find smORFs that overlap the regions of rRNAs, tRNAs, non-coding RNAs (ncRNAs), and pseudogenes as well. Interestingly, the only subgroups that are in the vicinity of ncRNAs are uORFs and dORFs, namely tORF 34630 and gORF 43305. The latter also overlaps a rRNA gene and, therefore, is found within a RNA cluster, and could have possible regulatory functions regarding these genes, as there are findings showing dORFs might enhance the translation of their canonical ORFs found within the same transcript ^20^.

After overlaying our results with the datasets from DeJesus et al. (2007) 21, which contain evidence for gene essentiality based on a saturated transposon mutagenesis technique, we found out that some smORFs are present in both annotated and unannotated genomic segments predicted to be essential or to affect bacterial growth (Supplementary Table 3). An important finding is that gORF 43305, which is the dORF found within a RNA cluster, is embedded by essential genomic segments - one that is present within the sequence of a rRNA, and another one that had yet to be annotated. Six of the novel smORFs are present in the 5’-UTR of known genes - two of which are classified as essential, three are responsible for a Growth Advantage (GA) when disrupted by a transposon, and one results in Growth Defect (GD) when disrupted. tORF 87698 is present in a GD region, and its transcript is differentially expressed upon nutrient starvation in the analyzed RNA-Seq datasets (Figure 3c), suggesting an important biological role for this smORF in bacterial growth. Furthermore, it is located nested in another reading frame of the transcript that encodes the two-component system sensor histidine kinase MtrB ^22^, which means it could extend a well known small regulatory system. gORF 250 is also essential and overlaps the segment of Rvnt02, which encodes a tRNA. Moreover, six smORFs are present in promoter segments that are either essential or GA, suggesting a possible regulatory mechanism for transcription.

**Figure 3.**
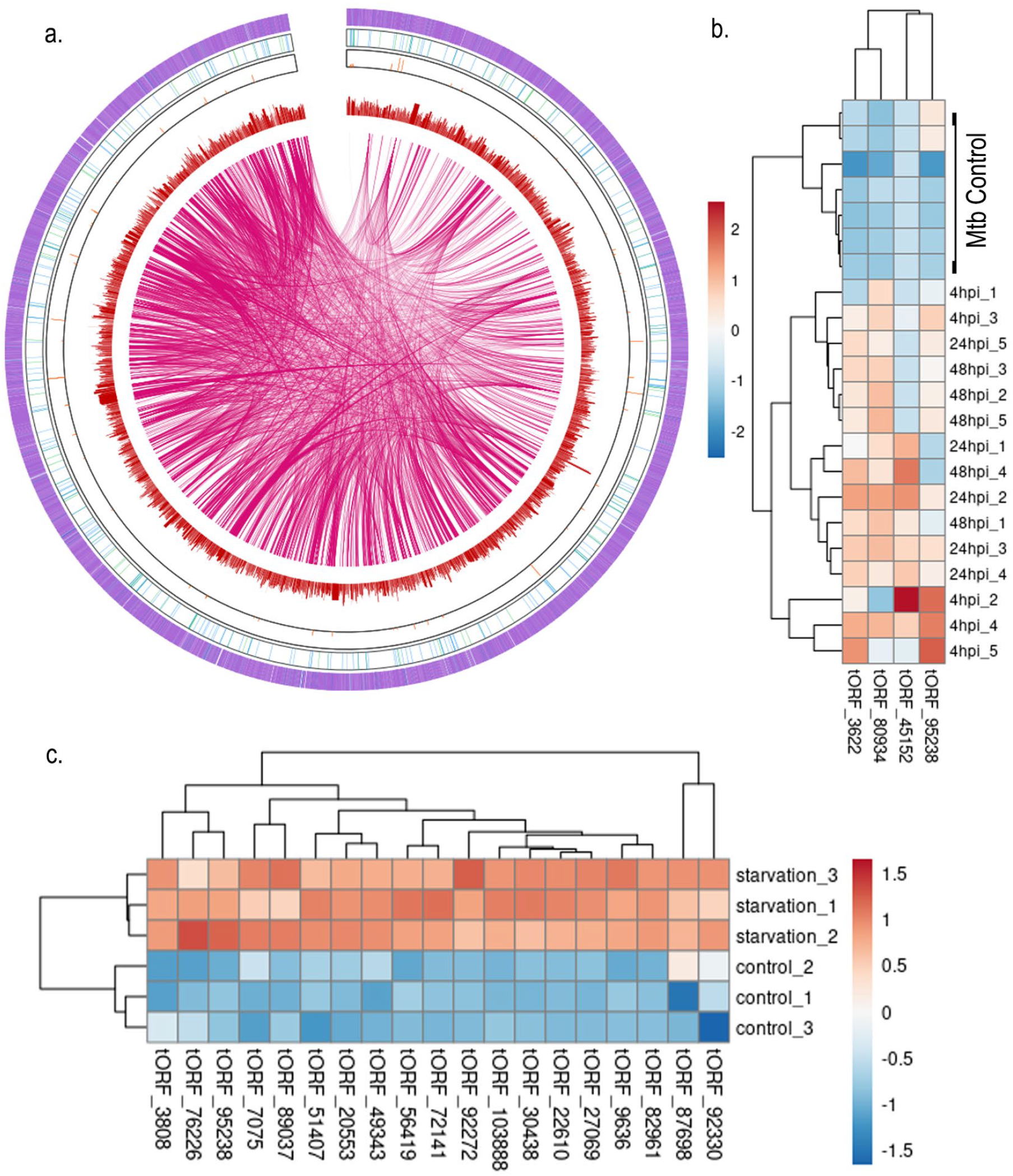
Genomic and transcriptomic landscape of novel smORFs. **a,** Circos plot representing the novel smORFs in a genomic context. The first ring, from edge to center, contains different purple-colored bars that represent the coordinates of annotated transcripts. In the second ring, smORFs are represented as blue bars. The third ring contains bars representing the number of homologous sequences in closely-related bacteria for smORFs in the same coordinates. The fourth section contains peaks representing the TPM values of both known and novel transcripts at that region. The center of the plot contains links connecting smORFs and annotated genes that are co-expressed after treatment with Isoniazid using public RNA-Seq datasets. **b,** Heatmap of differentially expressed SCTs during macrophage infection by *Mtb*. *Mtb* control refers to the bacterium grown in standard medium, without infecting the macrophages. **c,** Heatmap of differentially expressed SCTs upon nutrient starvation of *Mtb* cultures. SCT = smORF-containing Transcript, TPM = Transcript per Million.

### Most smORFs share operons with annotated genes

Aiming to further understand the biological roles of the novel smORFs and their encoded microproteins, we ran Rockhopper ^23^ on the same RNA-Seq dataset used for the transcriptome assembly to predict operons that contained already known genes. Afterwards, we mapped the coordinates of each novel smORF and checked which ones are located inside these predicted operons. This is useful when trying to infer functions for novel microproteins, as it is known that genes sharing the same operon usually have related biological roles ^24^. Hence, even though most microproteins possess only intrinsic disorder domains, it is still possible to contribute to their annotation by investigating their neighboring regions in the genome. Our findings show that the vast majority of both gORFs and tORFs (98.7%) share a mRNA with another known ORF, while only 5 tORFs were found in a novel transcript that does not contain any annotated CDS (figure 2 or 3 C). This trend seems to be common in eukaryotes as well, as more than 76% of the novel sequences found by Martinez et al ^11^ also seem to be found in known transcripts. Our data also shows that transcripts that would otherwise be predicted as monocistronic by Rockhopper also carry a smORF. More specifically, 217 gORFs and 66 tORFs were found within a transcript predicted to contain only one CDS. With the identification of these novel smORFs, we can classify these transcripts as polycistronic. Within the transcripts predicted to contain two or more annotated genes, we found 61 tORFs and 35 gORFs. Interestingly, tORF 72141 is present in the same region as the prophage-like element phiRv2. This prophage segment, along with phiRv1, is thought to be responsible for the low-level lysis that *Mtb* shows in culture ^25^. tORF 72141 shares this operon with many prophage genes, namely Rv2654c, Rv2655c, Rv2656c, Rv2657c, Rv2658c, and Rv2659c. As all the functions of these CDSs are related to the prophage biology, we can infer that this novel smORF is so as well.

Moreover, Houghton et al. has shown that Rv2660c, which is at that same locus, but in the opposite strand, is up-regulated during starvation and infection, and its transcript is expressed solely on strains that carry this prophage in their genome. The transcript that carries tORF 72141 is also found up-regulated upon starvation in the RNA-Seq dataset used in this study (Figure 3c), suggesting a positive correlation with their findings. Therefore, just like Rv2660c is described to be an useful biomarker during bacterial starvation, this novel smORF might also play a similar role ^26^.

**Figure 4.**
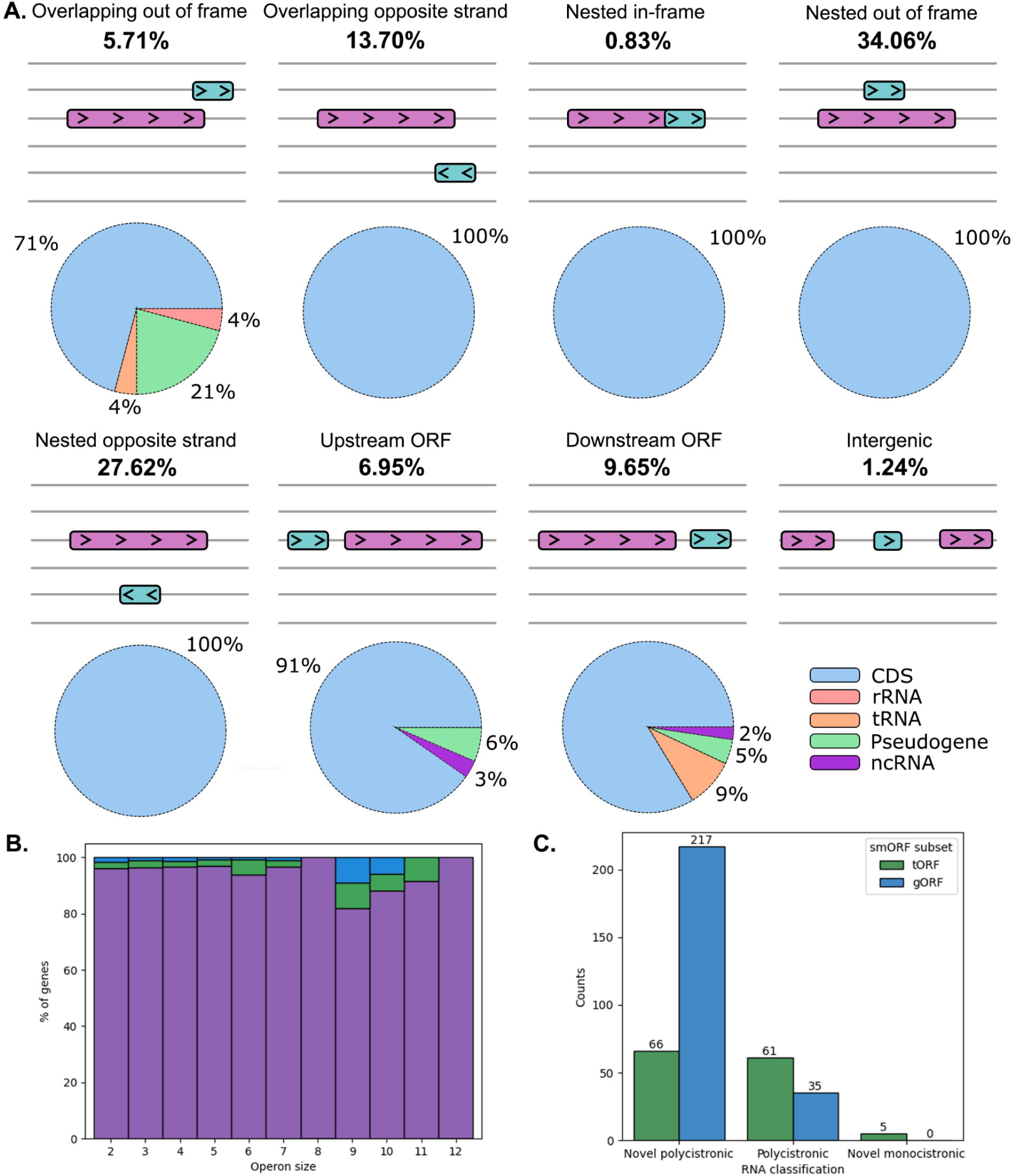
smORF neighborhood in the genome. **a,** classification of smORFs regarding their neighboring and overlapping regions in the genome. Each line represents a reading frame, purple rectangles represent annotated genes, cyan rectangles represent smORFs, and the arrows inside the boxes represent whether their coding sequences are in the forward or reverse strand. Pie charts on top of each classification show the proportion of gene types of overlapping or neighboring regions for each smORF. **b,** bar plot showing the % of annotated genes (purple), gORFs (blue) and tORFs (green) for each operon predicted by Rockhopper, which are grouped by the number of genes they encode. **c,** bar plot showing the class of operons that smORFs are a part of. Novel polycistronic means that these operons were predicted as monocistronic by Rockhopper and now include additional ORFs with the addition of the novel smORFs; polycistronic denotes operons that were already predicted to carry more than one gene and now carry smORFs as well; novel monocistronic denote mRNAs that are either novel or annotated as non-coding before, but now carry one smORF.

### SCTs are differentially expressed upon antibiotic treatment, bacterial starvation, and macrophage infection

To better integrate multi-omics datasets, we downloaded RNA-Seq reads from the study ^27^ deposited on Gene Expression Omnibus (GEO), accession number GSE166622, in which *Mtb* was exposed to different concentrations of a variety of antibiotics. Subsequently, we mapped the reads to a fasta file containing the nucleotide sequences of our transcriptome assembly and performed a differential expression analysis for the different antibiotics treatment, which included isoniazid, rifampicin and pyranizoic acid, and then filtered the results to include SCTs only. After exposure to isoniazid for 72h, tORF 92330 has the highest log2FoldChange among smORFs, and tORF 33916 has the highest log2FoldChange after exposure to rifampicin during 24h and 72h. Some SCTs are differentially expressed after exposure to more than one specific antibiotic, and some are exclusive to some time points across the conditions. For instance, there is only one SCT that is differentially expressed at 72h after exposure to all three antibiotics, which is the one that carries tORF_14178. This profile could present an important gene for late antibiotic response in all three treatments, just like tORF_3622 could act during the early response for rifampicin and isoniazid. tORFs 92330 and 49343 are both interesting for the antibiotic response overall as they are the two most prevalent after exposure to all three antibiotics in many time points. Complete data of differentially expressed tORFs upon exposure to antibiotics is available on Supplementary Table 4, and the groups in the UpSet plot and their corresponding smORFs are available on Supplementary Table 5. We also sought to characterize smORFs that are up or down regulated during bacterial starvation, in comparison to exponentially-grown cultures. To do so, we gathered the raw files from E-MTAB-1616 and performed a differential expression analysis at the transcript level. We found out that many SCTs are regulated in these conditions (Figure 3c, Supplementary Table 6), and tORF 87698 is present in a region whose deletion results in a Growth Defect. Therefore, it seems important for bacterial growth, especially in nutrient-lacking conditions. To gather insight into smORFs that are important for virulence, we also performed a differential expression analysis using data from dual RNA-Seq of THP-1 macrophages infected with *Mtb* (Figure 3b). We only found evidence for the differential expression of four smORFs, tORF 45152, 3622, 80934, and 95238, which were most expressive during the initial time points of the infection, at 4h and 24h post-infection, and thus could encode a subset of microproteins that play a role at the initial stages of macrophage infection. This low number of differentially expressed smORFs during infection could be due to conditional expression, where some genes would be expressed only during macrophage infection. In this case, simply mapping the reads to the novel genes in the transcriptome would not work, and a full proteogenomics analysis would have to be carried on, starting from the experimental steps.

### Co-expression of SCTs with known genes

Next, we sought to understand how these novel smORFs are connected to the rest of the transcriptome. To do so, we performed a co-expression analysis, which allows us to infer a correlation between the expression of two different genes based on similarities in their transcript expression level across multiple conditions. We then inferred a network based on the positive correlations we found in these datasets (Supplementary Table 7), for which we selected only the SCTs and their first neighbors in the network to represent in Figure 6.D. In contrast to looking directly at the smORFs sequences to infer their function, we decided to investigate the genes that are co-expressed with the SCTs in the network, as their functions are very likely similar or related to those in the same network, especially for those that act as a gene hub, defined as a gene with high connectivity and very likely to play a biological role. For those neighbors, we extracted their gene identifiers and performed a Gene Ontology (GO) overrepresentation analysis (Supplementary Table 8). As shown in Figure 6.D, the SCTs are part of larger networks, whose nodes represent genes with known biological roles. For instance, tORF 53851 is co-expressed with many genes whose biological processes are related to response to stimuli or chemicals. Also, some of these genes seem to partake in cellular processes, which is a common trend for many annotated genes in the landscape of these networks. tORF 92330, which has the highest log2FoldChange (3.22943411557331) among smORFs upon exposure to isoniazid for 72h and the second highest after exposure to pyranizoic acid (1.99608319151217) for 72h (figure 5a), is co-expressed with genes involved in cellular processes, response to stimuli, and protein binding (figure 6). Some of these genes encode proteins involved in bacterial pathogenicity, like ESAT-6-like protein EsxB ^28^, ESX-1 secretion-associated protein EspF ^29^, early secretory antigenic target EsxA ^28^, the antitoxin MazF6, and also PPE family protein PPE68, which is important for *Mtb* immunogenicity ^30^. Overexpression of MazF6 retards cell growth, and deletion of this gene makes the bacteria more sensitive to antibiotics ^31^. That coupled with tORF 92330 late response to both antibiotics could make the smORF attractive for studying antibiotic response, as well as tORF 103255, which has a similar co-expression profile and has the second highest log2FoldChange (2.70628414820118) after exposure to isoniazid for 72h.

**Figure 5.**
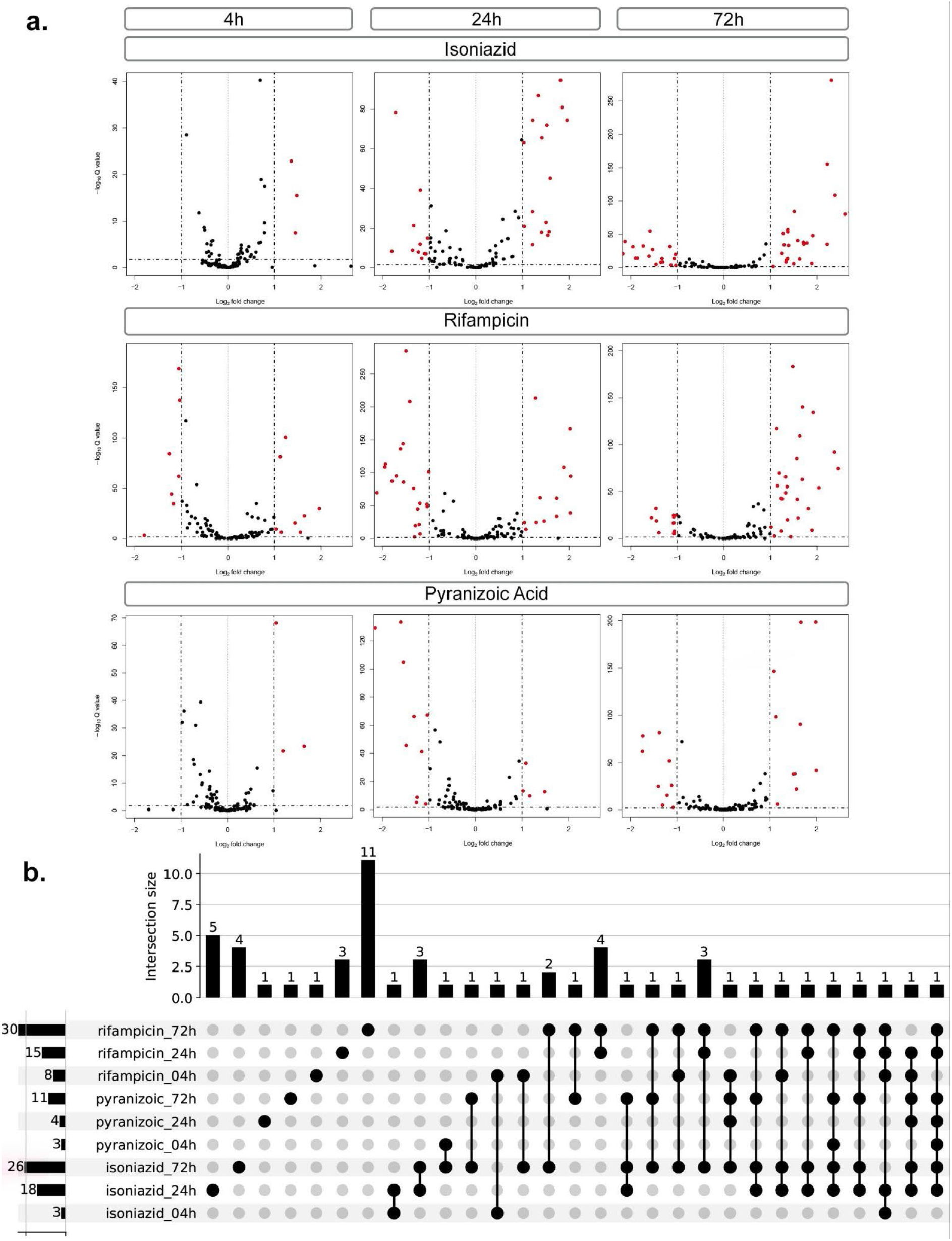
Differentially expressed SCTs upon antibiotic treatment. **a,** Volcano plots showing SCTs that are differentially expressed in one of the time points for each antibiotic compared to control. Red dots denote a SCT that has a padj < 0.05 and a log2FoldChange >1. **b,** UpSet plot containing groups that hold SCTs that are differentially expressed in one or more conditions. The dots on the center depict the groups in which the SCTs of that group are differentially expressed. Linked dots denote an intersection between one or more groups. Bars on the side show the number of differentially expressed SCTs for that treatment, and bars on the top show the number of SCTs in that intersect group. For instance, the two dots of isoniazid_24h and isoniazid_72h are linked, and there are 3 differentially expressed SCTs that are shared by those two treatments exclusively. (SCTS: smORF-Containing Transcripts, padj < adjusted p-value for multiple hypothesis testing).

### smORFs are widespread across the genome of closely-related bacteria and provide evidence for *de novo* gene birth events

Comparative genomics has been traditionally used to annotate functional genomic sequences, and is highly dependent on the principles described by Kimura (1983) ^32^. Said theory does not undermine Darwinian selection, but highlights another important aspect of molecular evolution, where most silent changes in the DNA and protein sequences are not adaptive in nature, but are in fact the result of random drift of neutral or nearly neutral mutants. Therefore, it is expected that most functional protein sequences share a degree of conservation across other species, especially when put into constraint as a result of purifying selection.

**Figure 6.**
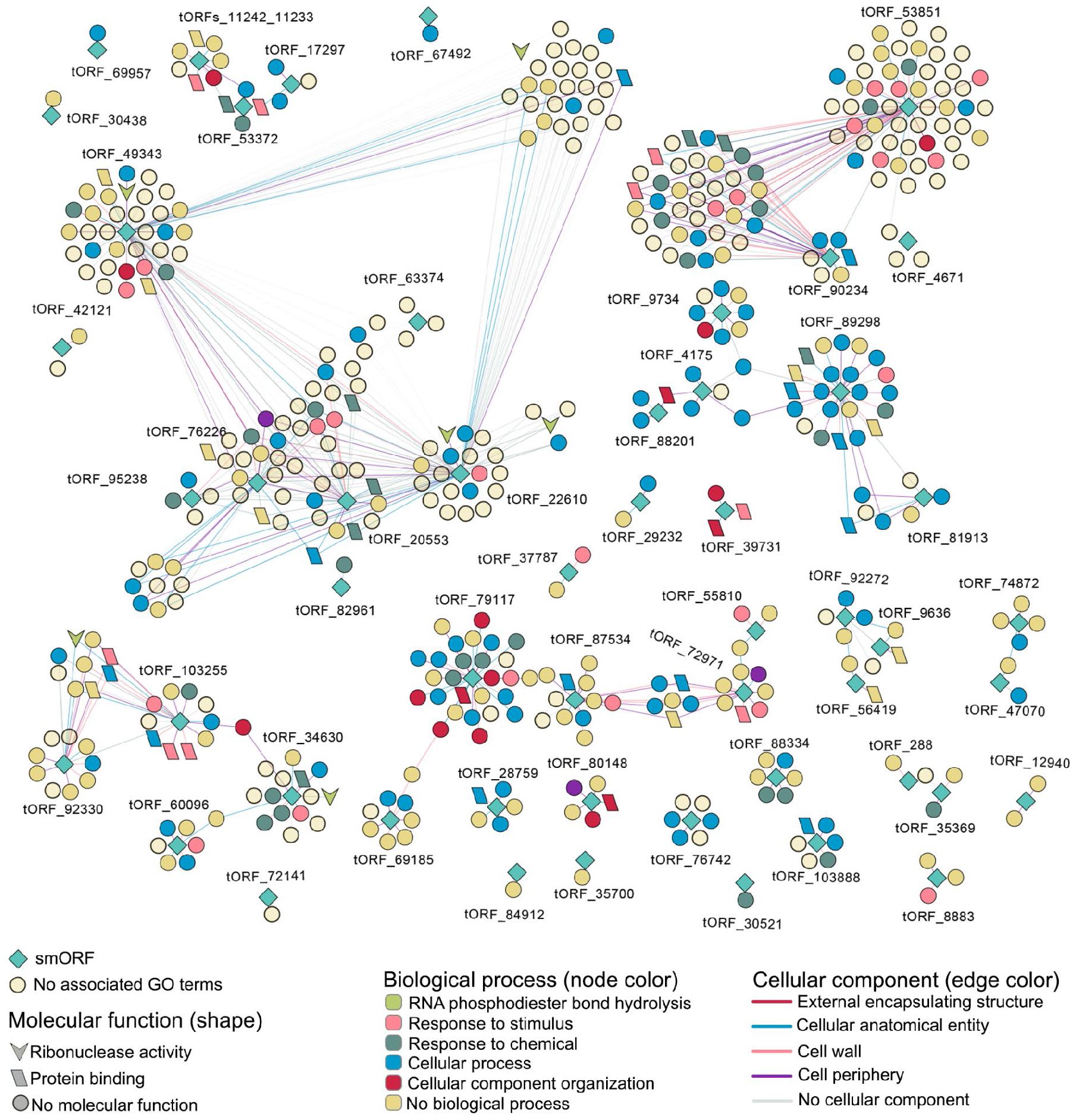
Co-expression networks of smORFs and annotated genes. The networks show smORFs and their first neighbors that are co-expressed in different RNA-Seq datasets used in this study. smORF nodes are represented as cyan diamonds, and the shapes of nodes representing annotated genes depend on their molecular function inferred from the GO overrepresentation analysis. Circles denote a node with no molecular function, downward arrows represent ribonuclease activity, and parallelograms denote a node with protein binding function. Biological processes are represented based on the node colors, and cellular components are illustrated based on the colors of the edge connecting different nodes. An annotated gene with no related GO terms is depicted as a node that is a beige-colored circle.

To gather insight into the evolution of the mycobacterial smORFome and identify which ones are conserved and thus more prone to be functional, we first identified which organisms from the Actinobacteria group have the potential to encode protein sequences with a certain degree of similarity to the novel microproteins (Supplementary Table 9). As shown in figure 7a, a wide array of bacteria from this phylum have at least one sequence homologous to one of the novel smORFs, whereas genera that are evolutionarily closer to *Mycobacterium* possess a higher number of homologs, as expected. Excluding *Mtb*, the *Mycobacterium* genus has homologs in the genome of its species for 309 smORFs, while the recently renamed *Mycolicibacterium* holds 108 homologs in total. Outside Mycobacteriaceae, both the genera *Nocardia* and *Rhodococcus* have homologous sequences for 69 smORFs spread across the genome of their species. If we probe deeper into the Mycobacteriaceae family (Fig 7.B), there is a clear evidence for the conservation of T1 and T2 smORFs in other genera, such as *M. canettii*, which possesses the highest number of homologous sequences. While some of these sequences share similarity with annotated sequences in these closely-related species, most of them have yet to be properly annotated in their genomes. The most conserved smORF is gORF 66553, while tORF 60096, gORF 68556, and tORF 87698 also share a high level of similarity to translated genomic segments. Interestingly, *M. smegmatis*, which is widely used as a model for studying tuberculosis, has no homologous sequences in its genome for any of the smORFs from T1 and T2, and only four for the T3 smORFs. A possible explanation for this would be the high discrepancy between the number of annotated genes in the two species, since the genome of *M. smegmatis* has 6938 genes against the 4173 from the much smaller genome of *Mtb* ^33^, many of which are responsible for its virulence, whereas *M. smegmatis* does not cause tuberculosis in humans. The tree also provides some insight about smORFs that are potentially important for virulence, as some of them are conserved in many pathogenic bacteria, such as *M. lepromatosis*, *M. leprae*, *M. canettii*, and *M. ulcerans*. A common trend in recent studies is the prevalence of smORFs coming from *de novo* gene birth events ^3–5^, where no conservation is found for the novel sequences, but there is still evidence for their translation. Together, this data suggests the importance of not undermining smORFs that are not conserved in other species, which led us to include all smORFs that are exclusive to *Mtb* and we could find evidence at the protein level for. Furthermore, genes that are unique to *Mtb* could represent a subset of smORFs that play an important role in its success as a pathogenic bacterium.

## Conclusion

Using our recently published pipeline, we were able to uncover a considerable amount of novel microproteins that had yet to be annotated in the genome and proteome of *Mtb.* By including an extensive amount of replicates, we classified the novel microproteins by degrees of confidence, and revealed the differences in the sequence composition of microproteins from different tiers. The annotation process of novel smORFs is usually made difficult by the absence of conserved protein domains and the prevalence of many intrinsic disorder regions, which was the pattern seen in our data, but also supported by many other studies. By overlaying multiple omics datasets, we evaluated the behavior of the novel smORFs at the transcript level across different conditions, which helped annotate these novel sequences. Moreover, using this multi-omics approach, we highlighted microproteins that are differentially regulated upon antibiotic treatment, which could be useful for studies focusing on resistance mechanisms and on the development of new anti-TB drugs. Many smORFs are also present in essential regions and are part of gene clusters with important biological functions. We believe our approaches can help establish a workflow to explore in-depth the function and cross-talk of any novel short sequences. Moreover, our analyses can help select which smORFs and microproteins should have higher priority when performing *in vitro* and *in vivo* experiments to further characterize the hidden microproteome of *Mtb*.

**Figure 7.**
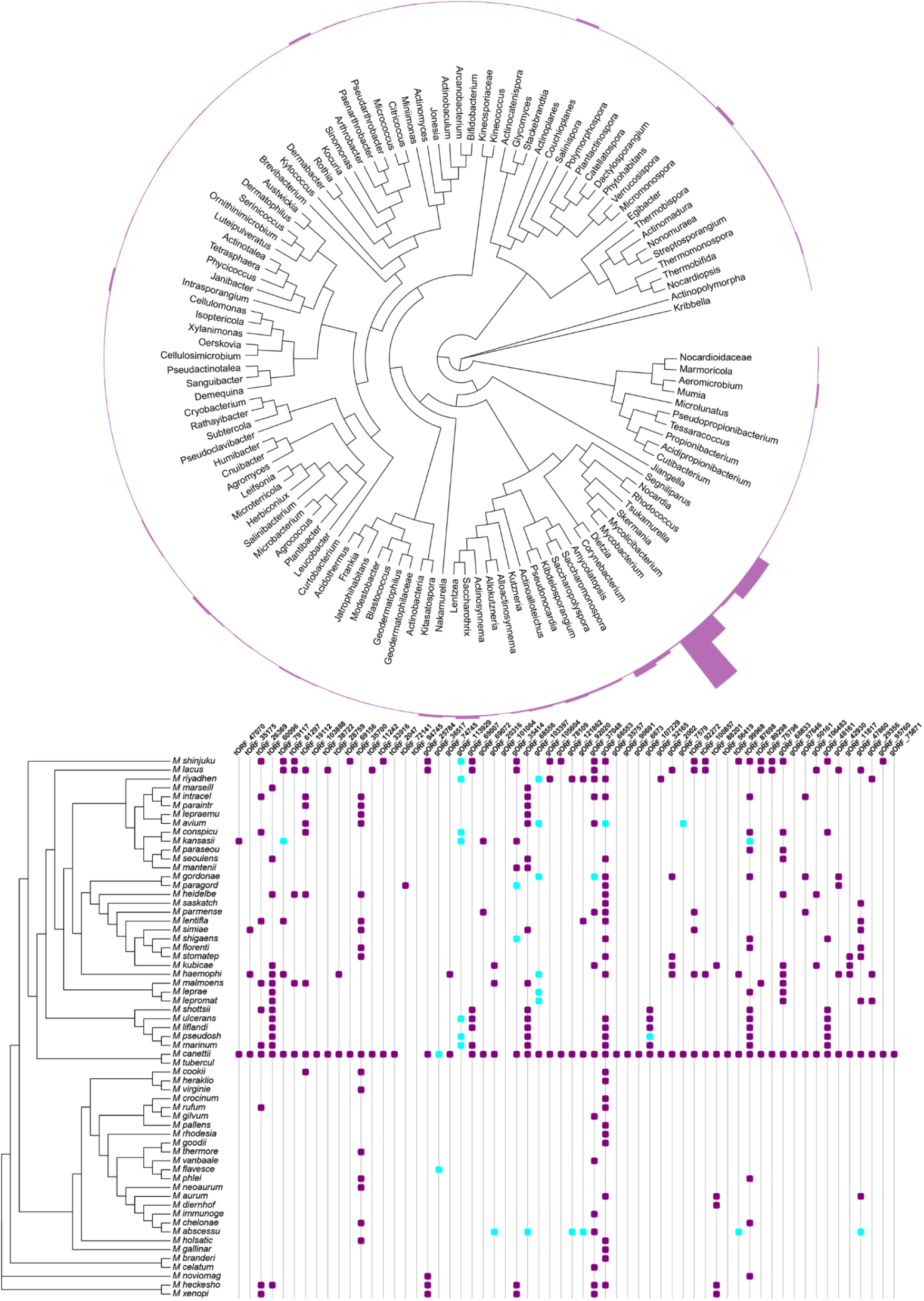
Phylogenetic analysis of novel microproteins. **a,** Unrooted phylogenetic tree inferred from high-quality 16S rRNA of Actinobacteria that have at least one tBlastn or Blastp hit for one of the novel microproteins. Bars around the tree depict the number of homologs in that genus. **b,** Matrix showing the level of conservation for the novel microproteins across Mycobacteria (*Mycobacterium* and *Mycolicibacterium).* Each microprotein in the matrix has a purple dot in its line for each unannotated homolog in the genome of that species (tBlastn hit) or a light blue dot for an annotated homolog (Blastp hit).

## Methods

### Data Acquisition

We downloaded the raw RNA-Seq reads in .fastq file for the Array Express studies E-MTAB-1616 ^34^ and E-MTAB-5287 ^35^, as well as from the study GSE166622 ^27^ from Gene Expression Omnibus (GEO). E-MTAB-1616 consists of 50 bp-long, single-end, strand-specific reads from a library of coding RNA from starved and exponentially grown *Mtb* cultures. E-MTAB-5287 reads are paired-end and 300 bp-long, obtained from macrophages infected with *Mtb.* The study GSE166622 from GEO consists of paired-end reads from *Mtb* exposed to a variety of antibiotics.

#### Protein extraction, in solution digestion and LC-MS/MS

We grew *Mtb* strains in 50 mL liquid medium (7H10 with OADC) at 37°C until an optical density (OD600) of 0.5-0.7 and washed cellular pellets twice using 10 mM Tris HCl pH 8.0. Cells were resuspended in 1 mL of the same buffer containing protease inhibitor cocktail (Promega, USA), and then transferred to 2 mL lysing matrix B tubes containing 0.1 mm diameter silica beads (MP Biomedicals, USA). We then disrupted the cells using a L-Beader 3 (Loccus, Brazil) at a speed setting of 4000 rpm, 10 cycles of 30 s each, cooling between cycles. The supernatants were filtered through 0.22 mM Millex Durapore (Millipore, USA). Some supernatants were filtered through 0.22 mM Millex Durapore (Millipore, USA), and triton X-114 (Sigma-Aldrich, USA) extraction was carried out to obtain detergent and aqueous fractions as described previously ^36^. Proteins were then precipitated with chloroform/methanol method ^37^. Protein pellets were resuspended in 100 mM tris HCl pH 7.0 (buffer A) containing 8 M urea, and digested with a protocol adapted from Klammer and MacCoss ^38^. Briefly, disulfide bonds were reduced in 5 mM dithiothreitol (DTT) for 20 min at 37 °C and then cysteines were alkylated in 25 mM iodoacetamide (IAM) for 20 min at room temperature in the dark. Urea was diluted to 2 M with 100 mM Tris HCl pH 7.0, trypsin was added at mass ratio of 1:100 (trypsin:protein) and the sample was incubated overnight at 37 °C. Formic acid was added (5% v/v, final concentration). Peptides were separated on an in-house made 20 cm revere-phase (5 µm ODSAQ C18, Yamamura Chemical Lab, Japan) using a nanoUPLC (nanoLC Ultra 1D plus, Eksigent, USA) connected to a LTQ-XL Orbitrap Discovery hybrid instrument (Thermo Fisher Scientific) through a nanoeletrospray ion source (Thermo Fisher Scientific). The flow rate was set to 300 nL min-1 in a 60, 120, 240 or 360 minutes reverse-phase gradient. The mass spectrometer was operated in a data-dependent mode, with full MS1 scan collected in the Orbitrap, with m/z range of 400-1600 at 30,000 resolution. The eight most abundant ions per scan were selected to CID MS2 in the ion trap.

### Reference-guided transcriptome assembly

Prior to the assembly, we performed the quality control of every RNA-Seq read used in the study using FastQC ^39^, to remove adapter contamination. We chose to run cutadapt and either perform a soft trimming for read quality when needed, or no trimming at all, as hard-trimming has been shown to negatively affect transcriptomics analysis ^40^. Afterwards, we used *assembly*, the first mode of µProteInS, to perform a reference-guided transcriptome assembly. Briefly, we fed to the pipeline the reference *Mtb* genome (NC 000962.3), a `GTF` annotation file, and the `.fastq` files containing the control RNA-Seq reads from the study E-MTAB-1616. Then, reads were aligned to the reference genome using HISAT2 ^41^ with the `--dta` flag. The resulting .`sam` alignment files were converted to sorted `.bam` files with the software Samtools ^42^. These sorted files were used as input for StringTie ^43^, along with the `GTF` annotation file. The minimum transcript length was set to 30 nucleotides. This was done for every replicate and, afterwards, we ran StringTie with the `--merge` flag, to generate a non-redundant, final set of transcripts that were stored in a `GTF` file.

### Generation of custom databases for proteomics

To generate a database containing all possible smORFs that are encoded by the transcriptome assembled in the previous step, we used the *database* mode of µProteInS. This part of the pipeline runs gffread ^44^ with the `-w` parameter to generate a fasta file containing the spliced-exon sequences that are extracted from the genome based on their coordinates in the `GTF` file that was generated during the transcriptome assembly. These sequences correspond to each transcript, and are translated to the three-reading frames using the specified start codons. We used ATG, ATT, GTG, and TTG, as these are known to be the most likely ones to initiate translation in mycobacteria ^21^. This mode also takes the same reference genome fasta file that was used during the transcriptome assembly and translates it to the six reading frames using the same set of start codons. Lastly, µProteInS appends the provided proteomes to the predicted databases and adds a tag to each entry from this proteome to signal that it is annotated. We used a concatenated proteome from NCBI RefSeq, Uniprot, and Mycobrowser to be truly comprehensive when excluding annotated proteins. To be even more restrictive, we performed a blastp search against NCBI RefSeq database for the H37Rv strain of *Mtb,* and removed microproteins that had 100% identity and query coverage, i.e. a perfect match. Sequences added to their database after the search may be similar to the novel smORFs.

### Bottom-up label free proteomics

We fed µProteInS third mode, *ms*, with 680 MS experiments in the `mzML` format. The conditions themselves are not taken into account here, and the sheer number of experiments are used to primarily increase the probability of an identification, given the stochastic nature of a label-free proteomics experiment. The pipeline uses MS-GF+ ^45^ as the search engine. We specified the following parameters: precursor mass tolerance of 20 ppm, isotope error range between 0 and 1, CID as fragmentation method, 2 tolerable termini (includes semi-tryptic peptides), peptide length between 6 and 40 amino acids, LCQ/LTQ as instrument, and Cysteine carbamidomethylation as a fixed modification. We performed the post-processing of the search using *postms*, the fourth mode of µProteInS, which uses Percolator ^46^ to assess the FDR. Throughout the pipeline’s workflow, FDR cutoffs of 1% at both the PSM and Peptide levels are applied, and any peptide that matches an annotated protein that is present in the reference annotation files is removed from the subsequent analyses. FDR assessment was done in chunks of 10 files, given Percolator limitations when handling memory for too many files. Since this is not a quantitative proteomics analysis, the replicates are used to identify smORFs in a qualitative and binary approach, rather than using the spectral counts coming from different conditions to infer differences in protein expression. To quantify smORFs, analyses were done at the transcript level using RNA-Seq data. To confirm that no annotated proteins were mistakenly identified as novel, we performed a Blastp search against NCBI RefSeq database for *Mtb* H37Rv strain, with an E-value cutoff of 0.01, optimized search for short sequences, and the filter for low complexity regions disabled. We then proceeded to remove any novel microprotein that had at least a hit with both 100 % identity and query coverage, i.e. a perfect match.

#### Definition of start codon priority and RBS detection

RBS were predicted using Free2Bind ^17^, and the most likely smORF for a stop codon with multiple alternative start codons was selected based on µProteInS selection criteria. First, priority is given to smORFs starting with an ATG. If there are multiple smORFs with the canonical start codon, they are sorted by length and the longest one is given priority. Next, they are sorted by the amount of free energy resulting from the binding to the upstream region of that start codon to the 3’ end of the 16S rRNA of *Mtb,* where lower free energy levels represent a RBS with stronger binding capacity, thus more likely to be translated. Finally, the pipeline checks if a Mass Spectrometry peptide is matching one microprotein coming from a smORF with an alternative start that is more upstream and not matching the other short smORFs, even if they were given priority before.

### Prediction of high-confidence PSMs

To check for strongly reliable evidence of microproteins encoded by smORFs, we employed the fifth and last mode of µProteInS to predict spectra of high quality. Briefly, this mode applies a pre-trained random forest model, which was trained on manually inspected and curated Mass Spectrometry data, to identify features of a PSM that are related to those of a high-quality PSM. Each PSM is then assigned a classification based on the machine learning prediction, and a final result file is generated, containing only HC smORFs. We then combined the predicted results with the results coming from the *postms* mode to classify them into Tiers. To do so, we used a custom in-house python script to check in how many replicates each one of these smORFs appeared, and whether they were classified as HC or not.

### Function prediction

To predict conserved protein domains, we used the microprotein sequences from T1 to T3 as input for InterPro ^47,48^ and CDD ^49^. To predict cellular localization, we used Psortb ^50^. To predict transmembrane and signal peptide domains, we ran TMHMM ^51^, Phobius ^52^ and SignalP ^53^. Afterwards, to infer the Gene Ontology from the sequence, we ran CELLO ^54^ and DeepGoWeb ^55^.

### Differential expression analysis

We used the same assembly generated with the first mode of µProteInS to look for SCTs that might be differentially expressed in multiple conditions. First, we renamed each SCT in the fasta file to include the names of each tORF it carries, to improve visualization and interpretation of the results. The following steps were done for all samples from the downloaded RNA-Seq studies. E-MTAB-5287 was the only one that required an additional step, where we aligned the reads to the human genome (GRCh37) and kept only reads that did not map to it, which should be coming from the transcriptome of *Mtb*. We then indexed the *Mtb* transcriptome using the genome as decoy, as described in Salmon documentation, and ran Salmon in `quant` mode with the flag `--validateMappings`^56^. We created a file containing the read counts from each one of the output `quant.sf` files from Salmon and, prior to normalization, excluded low read counts that matched the following criteria: genes with less than 3 samples with normalized counts greater or equal to 5. Then, we used DESeq2 ^57^ to perform the differential gene expression analysis at the transcript level. After normalization, we excluded transcripts that did not carry a smORF from Tiers 1 to 3. Afterwards, we applied a cutoff for an adjusted p-value (padj) of 0.05, and for a Log2FoldChange of 1. Next, we applied a regularized log transformation to the DESeq2 results, and plotted them into a heatmap using the package pheatmap (https://www.rdocumentation.org/packages/pheatmap/versions/1.0.12/topics/pheatmap) and into volcano plots.

### Co-expression analysis

Using the same file containing raw counts from the differential expression analysis, we used the EBSeq R package ^58^ to get a median normalized matrix. Then, for each possible combination, we performed a pairwise Pearson correlation analysis using the R package Psych ^59^. Afterwards, we selected gene pairs with a padj < 0.05 and an absolute correlation >= 0.9 and used them as input for the igraph ^60^ package, to calculate metrics such as degree and betweenness centrality for each gene pair. To put it simply, degree is calculated based upon the number of connections a node makes, and betweenness centrality measures how many times a specific node is used as a path to connect two or more other nodes. In the network, each node corresponds to a gene, be it annotated or novel, and the edges correspond to a correlation between the expression level of two different nodes across different conditions. We then visualized the network and node files in Cytoscape ^61^, where we selected an organic layout and manually adjusted the networks to improve visualization. Then, to assess information about the Gene Ontology (GO) of annotated genes that are co-expressed with one or more smORFs, we extracted their gene terms and performed a GO overrepresentation analysis using the Gene Ontology (http://geneontology.org/) database. As the background gene list to infer p-values, we used all *Mtb* genes in the database, and filtered out of the results any hit with a p-value > 0.05. Then, we customized the co-expression networks based on the first GO terms in the list associated with each gene, which have the highest fold enrichment. For biological processes, we associated each GO term with a specific color, and then colored the network nodes for each gene based on its GO term color. For cellular component, we associated each edge color with a different term. Lastly, for GO terms related to molecular function, we defined a node shape for each different term.

### Conservation analyses of smORFs in closely-related species within the Actinobacteriaceae family

To build the unrooted phylogenetic tree and its matrix containing all homologous sequences, we performed three sets of Blast searches. To look for annotated conserved sequences, we ran Blastp using the microproteins sequences encoded by smORFS from T1 to T3 as query, and searched the non-redundant NCBI database twice - once restricting the search to Actinobacteria (taxid:201174) and excluding Mycobacteriales (taxid:85007), and then restricting the search to Mycobacteria and excluding *Mtb* (taxid:1773). We set the E-value cutoff to 0.001 and filtered out hits with a BitScore < 50. As the low complexity filter sometimes negatively affects the number of hits for smORFs, undermining the number of conserved sequences ^5^, we turned it off for the searches. To check for unannotated conserved sequences, we ran tBlastN using the same parameters and database restrictions. Next, we extracted the species from the entries of each blast hit that had a complete genome and downloaded their 16S rRNA sequences from the Silva database ^62^, which collects high-quality ribosomal sequences for many different organisms. Subsequently, we performed a multiple sequence alignment using MAFFT ^63^ and used this as input for PhyML ^64^ to reconstruct the unrooted phylogenetic tree from the 16S rRNA sequences. To generate the matrix containing conservation information for each species in the tree (Fig 7b) and the bars for the Actinobacteriaceae tree (Fig 7a), we used EvolView ^65^. Shortly, for each smORF that had a hit in the blastP search, we added a purple-colored box to the matrix, and an orange-colored box for each smORF that did not have a blastP hit, but had a tBlastN hit for the species of the corresponding leaf in the tree. For the bars in Fig 7a, we added the absolute numbers of smORFs homologs in the genomes of each species. Full genera for the species in the tree are provided in Supplementary Table 10.

### Statistical analyses

For each comparison that required statistical inference, we used the Python packages SciPy (https://scipy.org/) and Statannotations (https://github.com/trevismd/statannotations) to perform the analyses. First, we checked for a Gaussian distribution for each group using the Shapiro-Wilk test. For samples whose tests rejected H0 and thus had a non-parametric distribution, we used the Kruskal-Wallis test for pairwise comparison of independent variables between 2 or more groups. For samples that failed to reject H0 and followed a parametric distribution, we used paired T-tests. Each comparison was corrected afterwards for multiple hypothesis testing using the Bonferroni correction.

## Acknowledgements

The authors acknowledge the High-Performance Computing Laboratory of the Pontifical Catholic University of Rio Grande do Sul (LAD-IDEIA/PUCRS, Brazil) for providing support and technological resources. This work was supported by CNPq/FAPERGS/CAPES/BNDES (INCT-TB) [421703-2017-2/17-1265-8/14.2.0914.1]. C.V.B. [310344/2016-6], P.M. [305203/2018-5] and L.A.B. [520182/99-5] are research career awardees of the National Council for Scientific and Technological Development of Brazil (CNPq). This study was financed in part by the Coordenação de Aperfeiçoamento de Pessoal de Nível Superior— Brasil (CAPES)—Finance Code 001.

## Author contributions

EVS, PFD, AMP, CVB, AS: conception and design of the work; EVS, PFD, AMP: data acquisition; EVS, PFD, ACM, AS, CVB: analysis and interpretation of data; PM, LAB, AS, CVB: supervision, project administration, funding acquisition; EVS: writing-original draft; EVS, ACM, CVB, AS: writing-review and editing.

## Competing interests

The authors declare no competing interests.

## Materials & Correspondence

Correspondence and material requests should be addressed to CVB.

## Notes

### Competing Interest Statement

The authors have declared no competing interest.

### Summary of Updates

Supplemental Tables 1 and 6 updated.

## References

1. Orr, M. W., Mao, Y., Storz, G. & Qian, S. B. Alternative ORFs and small ORFs: shedding light on the dark proteome. Nucleic Acids Res. (2020) doi:10.1093/nar/gkz734.

2. Basrai, M. A., Hieter, P. & Boeke, J. D. Small open reading frames: beautiful needles in the haystack. Genome Res. 7, 768–771 (1997).

3. Schmitz, J. F. & Bornberg-Bauer, E. Fact or fiction: updates on how protein-coding genes might emerge de novo from previously non-coding DNA. F1000Research 6, (2017).

4. Ruiz-Orera, J., Verdaguer-Grau, P., Villanueva-Cañas, J. L., Messeguer, X. & Albà, M. Translation of neutrally evolving peptides provides a basis for de novo gene evolution. Nat. Ecol. Evol. 2, 890–896 (2018).

5. Fesenko, I. et al. A vast pool of lineage-specific microproteins encoded by long non-coding RNAs in plants. Nucleic Acids Res. 49, 10328–10346 (2021).

6. Guerra-Almeida, D. & Nunes-da-Fonseca, R. Small open reading frames: how important are they for molecular evolution? Front. Genet. 11, 574737 (2020).

7. Zhu, Y. et al. Discovery of coding regions in the human genome by integrated proteogenomics analysis workflow. Nat. Commun. 9, 1–14 (2018).

8. Fuchs, S. et al. Towards the characterization of the hidden world of small proteins in Staphylococcus aureus, a proteogenomics approach. PLoS Genet. 17, e1009585 (2021).

9. Potgieter, M. G. et al. Proteogenomic analysis of mycobacterium smegmatis using high resolution mass spectrometry. Front. Microbiol. 7, 427 (2016).

10. Smith, C. et al. Pervasive translation in Mycobacterium tuberculosis. Elife 11, e73980 (2022).

11. Martinez, T. F. et al. Accurate annotation of human protein-coding small open reading frames. Nat. Chem. Biol. 16, 458–468 (2020).

12. Chothani, S. P. et al. A high-resolution map of human RNA translation. Mol. Cell 82, 2885–2899 (2022).

13. Xiao, Z. et al. De novo annotation and characterization of the translatome with ribosome profiling data. Nucleic Acids Res. 46, e61–e61 (2018).

14. de Souza, E. V. et al. μProteInS—a proteogenomics pipeline for finding novel bacterial microproteins encoded by small ORFs. Bioinformatics 38, 2612–2614 (2022).

15. Venter, E., Smith, R. D. & Payne, S. H. Proteogenomic analysis of bacteria and archaea: a 46 organism case study. PloS One 6, e27587 (2011).

16. Tsiatsiani, L. & Heck, A. J. Proteomics beyond trypsin. FEBS J. 282, 2612–2626 (2015).

17. Starmer, J., Stomp, A., Vouk, M. & Bitzer, D. Predicting Shine–Dalgarno sequence locations exposes genome annotation errors. PLoS Comput. Biol. 2, e57 (2006).

18. Guerra-Almeida, D., Tschoeke, D. A. & Nunes-da-Fonseca, R. Understanding small ORF diversity through a comprehensive transcription feature classification. DNA Res. 28, dsab007 (2021).

19. Wright, B. W., Molloy, M. P. & Jaschke, P. R. Overlapping genes in natural and engineered genomes. Nat. Rev. Genet. 1–15 (2021).

20. Wu, Q. et al. Translation of small downstream ORFs enhances translation of canonical main open reading frames. EMBO J. 39, (2020).

21. DeJesus, M. A., Sacchettini, J. C. & Ioerger, T. R. Reannotation of translational start sites in the genome of Mycobacterium tuberculosis. Tuberculosis 93, 18–25 (2013).

22. Zahrt, T. C. & Deretic, V. An essential two-component signal transduction system in Mycobacterium tuberculosis. J. Bacteriol. 182, 3832–3838 (2000).

23. Tjaden, B. De novo assembly of bacterial transcriptomes from RNA-seq data. Genome Biol. 16, 1–10 (2015).

24. Osbourn, A. E. & Field, B. Operons. Cell. Mol. Life Sci. 66, 3755–3775 (2009).

25. Cole, S. T. et al. Deciphering the biology of mycobacterium tuberculosis from the complete genome sequence. Nature (1998) doi:10.1038/31159.

26. Houghton, J. et al. A small RNA encoded in the Rv2660c locus of Mycobacterium tuberculosis is induced during starvation and infection. PloS One 8, e80047 (2013).

27. Srinivas, V. et al. Transcriptome signature of cell viability predicts drug response and drug interaction in Mycobacterium tuberculosis. Cell Rep. Methods 1, 100123 (2021).

28. Burts, M. L., Williams, W. A., DeBord, K. & Missiakas, D. M. EsxA and EsxB are secreted by an ESAT-6-like system that is required for the pathogenesis of Staphylococcus aureus infections. Proc. Natl. Acad. Sci. 102, 1169–1174 (2005).

29. Bottai, D. et al. ESAT-6 secretion-independent impact of ESX-1 genes espF and espG 1 on virulence of Mycobacterium tuberculosis. J. Infect. Dis. 203, 1155–1164 (2011).

30. Demangel, C. et al. Cell envelope protein PPE68 contributes to Mycobacterium tuberculosis RD1 immunogenicity independently of a 10-kilodalton culture filtrate protein and ESAT-6. Infect. Immun. 72, 2170–2176 (2004).

31. Kumari, K. & Sarma, S. P. Structural and mutational analysis of MazE6-operator DNA complex provide insights into autoregulation of toxin-antitoxin systems. Commun. Biol. 5, 1–14 (2022).

32. Kimura, M. The Neutral Theory of Molecular Evolution Cambridge University. NY Camb. (1983).

33. Kapopoulou, A., Lew, J. M. & Cole, S. T. The MycoBrowser portal: a comprehensive and manually annotated resource for mycobacterial genomes. Tuberculosis 91, 8–13 (2011).

34. Cortes, T. et al. Resource Genome-wide Mapping of Transcriptional Start Sites Defines an Extensive Leaderless Transcriptome in Mycobacterium tuberculosis. CellReports 5, 1121–1131 (2013).

35. Zimmermann, M. et al. Integration of Metabolomics and Transcriptomics Reveals a Complex Diet of Mycobacterium tuberculosis during Early Macrophage Infection. mSystems 2, (2017).

36. Målen, H., Pathak, S., Søfteland, T., De Souza, G. A. & Wiker, H. G. Definition of novel cell envelope associated proteins in Triton X-114 extracts of Mycobacterium tuberculosis H37Rv. BMC Microbiol. 10, 1–11 (2010).

37. Wessel, D. & Flügge, U. A method for the quantitative recovery of protein in dilute solution in the presence of detergents and lipids. Anal. Biochem. 138, 141–143 (1984).

38. Klammer, A. A. & MacCoss, M. J. Effects of modified digestion schemes on the identification of proteins from complex mixtures. J. Proteome Res. 5, 695–700 (2006).

39. Andrews, S. FastQC.

40. Williams, C. R., Baccarella, A., Parrish, J. Z. & Kim, C. C. Trimming of sequence reads alters RNA-Seq gene expression estimates. BMC Bioinformatics 17, 103 (2016).

41. Kim, D., Paggi, J. M., Park, C., Bennett, C. & Salzberg, S. L. Graph-based genome alignment and genotyping with HISAT2 and HISAT-genotype. Nat. Biotechnol. 37, 907– 915 (2019).

42. Li, H. et al. The sequence alignment/map format and SAMtools. Bioinformatics 25, 2078–2079 (2009).

43. Pertea, M. et al. StringTie enables improved reconstruction of a transcriptome from RNA-seq reads. Nat Biotechnol 33, 290–295 (2015).

44. Pertea, G. & Pertea, M. GFF utilities: GffRead and GffCompare. F1000Research 9, (2020).

45. Kim, S. & Pevzner, P. A. MS-GF+ makes progress towards a universal database search tool for proteomics. Nat. Commun. (2014) doi:10.1038/ncomms6277.

46. Käll, L., Canterbury, J. D., Weston, J., Noble, W. S. & MacCoss, M. J. Semi-supervised learning for peptide identification from shotgun proteomics datasets. Nat. Methods 4, 923–925 (2007).

47. Hunter, S. et al. InterPro: the integrative protein signature database. Nucleic Acids Res. 37, D211–D215 (2009).

48. Quevillon, E. et al. InterProScan: Protein domains identifier. Nucleic Acids Res. (2005) doi:10.1093/nar/gki442.

49. Marchler-Bauer, A. et al. CDD: NCBI’s conserved domain database. Nucleic Acids Res. 43, D222–D226 (2015).

50. Rey, S. et al. PSORTdb: a protein subcellular localization database for bacteria. Nucleic Acids Res. 33, D164–D168 (2005).

51. Krogh, A., Larsson, B., Von Heijne, G. & Sonnhammer, E. L. Predicting transmembrane protein topology with a hidden Markov model: application to complete genomes. J. Mol. Biol. 305, 567–580 (2001).

52. Käll, L., Krogh, A. & Sonnhammer, E. L. Advantages of combined transmembrane topology and signal peptide prediction—the Phobius web server. Nucleic Acids Res. 35, W429–W432 (2007).

53. Teufel, F. et al. SignalP 6.0 predicts all five types of signal peptides using protein language models. Nat. Biotechnol. 1–3 (2022).

54. Yu, C.-S. et al. CELLO2GO: a web server for protein subCELlular LOcalization prediction with functional gene ontology annotation. PloS One 9, e99368 (2014).

55. Kulmanov, M., Zhapa-Camacho, F. & Hoehndorf, R. DeepGOWeb: fast and accurate protein function prediction on the (Semantic) Web. Nucleic Acids Res. 49, W140–W146 (2021).

56. Patro, R., Duggal, G., Love, M. I., Irizarry, R. A. & Kingsford, C. Salmon provides fast and bias-aware quantification of transcript expression. Nat. Methods 14, 417–419 (2017).

57. Love, M. I., Huber, W. & Anders, S. Moderated estimation of fold change and dispersion for RNA-seq data with DESeq2. Genome Biol. 15, 1–21 (2014).

58. Leng, N. et al. EBSeq: an empirical Bayes hierarchical model for inference in RNA-seq experiments. Bioinformatics 29, 1035–1043 (2013).

59. Revelle, W. & Revelle, M. W. Package ‘psych’. Compr. R Arch. Netw. 337, 338 (2015).

60. Csardi, G., Nepusz, T., & others. The igraph software package for complex network research. InterJournal Complex Syst. 1695, 1–9 (2006).

61. Shannon, P. et al. Cytoscape: a software environment for integrated models of biomolecular interaction networks. Genome Res. 13, 2498–2504 (2003).

62. Quast, C. et al. The SILVA ribosomal RNA gene database project: improved data processing and web-based tools. Nucleic Acids Res. 41, D590–D596 (2012).

63. Katoh, K. & Toh, H. Recent developments in the MAFFT multiple sequence alignment program. Brief. Bioinform. 9, 286–298 (2008).

64. Guindon, S., Delsuc, F., Dufayard, J.-F. & Gascuel, O. Estimating maximum likelihood phylogenies with PhyML. in Bioinformatics for DNA sequence analysis 113– 137 (Springer, 2009).

65. Zhang, H., Gao, S., Lercher, M. J., Hu, S. & Chen, W.-H. EvolView, an online tool for visualizing, annotating and managing phylogenetic trees. Nucleic Acids Res. 40, W569– W572 (2012).

