## Supplementary Tables 2,3,5,8,9, and 10 for "Large-scale proteogenomics characterization of the *Mycobacterium tuberculosis* hidden microproteome"

### Supplementary Data

Supplementary Table 1. Conserved protein domains within novel microproteins. Available as a separate Excel file (see Supplementary Materials).

Supplementary Table 2. Overlapping regions in the genome. Each row represents a different gene that is overlapped in the genome by one of the novel smORFs. Surrounding gene ids include genes in the vicinity of the smORF, up to 500 bp.

| novel_ORF | Overlapping gene id | Overlapping gene name | Surrounding gene Ids | Surrounding gene names |
|---|---|---|---|---|
| gORF_206 | gene-Rv0006 | gyrA | | |
| gORF_206 | gene-Rv0006 | gyrA | | |
| gORF_250 | gene-Rvnt02 | alaT | gene-Rv0007 | |
| gORF_250 | gene-Rvnt02 | alaT | gene-Rv0007 | |
| gORF_121862 | gene-Rv0009 | ppiA | gene-Rv0008c | |
| gORF_121862 | gene-Rv0009 | ppiA | gene-Rv0008c | |
| gORF_422 | gene-Rv0014c | pknB | | |
| gORF_422 | gene-Rv0014c | pknB | | |
| gORF_806 | gene-Rv0021c | | gene-Rv0020c | fhaA |
| gORF_806 | gene-Rv0021c | | gene-Rv0020c | fhaA |
| gORF_1601 | gene-Rv0048c | | gene-Rv0047c | |
| gORF_1601 | gene-Rv0048c | | gene-Rv0047c | |
| gORF_120753 | gene-Rv0050 | ponA1 | | |
| gORF_120753 | gene-Rv0050 | ponA1 | | |
| gORF_119201 | gene-Rv0099 | fadD10 | | |
| gORF_119201 | gene-Rv0099 | fadD10 | | |
| gORF_3585 | gene-Rv0107c | ctpI | gene-Rv0106 | |
| gORF_3585 | gene-Rv0107c | ctpI | gene-Rv0106 | |
| gORF_118619 | gene-Rv0108c | | gene-Rv0107c | ctpI |
| gORF_118619 | gene-Rv0108c | | gene-Rv0107c | ctpI |
| gORF_118158 | gene-Rv0121c | | gene-Rv0120c | fusA2 |
| gORF_118158 | gene-Rv0121c | | gene-Rv0120c | fusA2 |
| gORF_4318 | gene-Rv0124 | PE_PGRS2 | | |
| gORF_4318 | gene-Rv0124 | PE_PGRS2 | | |
| gORF_117472 | gene-Rv0147 | | | |
| gORF_117472 | gene-Rv0147 | | | |
| gORF_5065 | gene-Rv0151c | PE1 | | |
| gORF_5065 | gene-Rv0151c | PE1 | | |
| gORF_117009 | gene-Rv0162c | adhE1 | | |
| gORF_117009 | gene-Rv0162c | adhE1 | | |
| gORF_117009 | gene-Rv0163 | | | |
| gORF_117009 | gene-Rv0163 | | | |
| gORF_6673 | gene-Rv0197 | | gene-Rv0196 | |
| gORF_6673 | gene-Rv0197 | | gene-Rv0196 | |
| gORF_115364 | gene-Rv0212c | nadR | | |

| | | | | |
|---|---|---|---|---|
| gORF_115364 | gene-Rv0212c | nadR | | |
| gORF_115364 | gene-Rv0213c | | | |
| gORF_115364 | gene-Rv0213c | | | |
| gORF_7303 | gene-Rv0213c | | gene-Rv0212c | nadR |
| gORF_7303 | gene-Rv0213c | | gene-Rv0212c | nadR |
| gORF_7760 | gene-Rv0226c | | | |
| gORF_7760 | gene-Rv0226c | | | |
| gORF_114282 | gene-Rv0249c | | gene-Rv0248c | |
| gORF_114282 | gene-Rv0249c | | gene-Rv0248c | |
| gORF_113520 | gene-Rv0278c | | gene-Rv0277c | vapC25 |
| gORF_113520 | gene-Rv0278c | | gene-Rv0277c | vapC25 |
| gORF_113073 | gene-Rv0286 | PPE4 | | |
| gORF_113073 | gene-Rv0286 | PPE4 | | |
| gORF_11107 | gene-Rv0309 | | | |
| gORF_11107 | gene-Rv0309 | | | |
| gORF_111934 | gene-Rv0336 | | gene-Rv0335c | PE6 |
| gORF_111934 | gene-Rv0336 | | gene-Rv0335c | PE6 |
| gORF_111269 | gene-Rv0355c | PPE8 | | |
| gORF_111269 | gene-Rv0355c | PPE8 | | |
| gORF_111295 | gene-Rv0355c | PPE8 | | |
| gORF_111295 | gene-Rv0355c | PPE8 | | |
| gORF_15589 | gene-Rv0429c | def | gene-Rv0428c | |
| gORF_15589 | gene-Rv0429c | def | gene-Rv0428c | |
| gORF_109100 | gene-Rv0439c | | | |
| gORF_109100 | gene-Rv0439c | | | |
| gORF_107153 | gene-Rv0509 | hemA | | |
| gORF_107153 | gene-Rv0509 | hemA | | |
| gORF_18752 | gene-Rv0533c | fabH | | |
| gORF_18752 | gene-Rv0533c | fabH | | |
| gORF_106483 | gene-Rv0535 | | | |
| gORF_106483 | gene-Rv0535 | | | |
| gORF_106292 | gene-Rv0545c | pitA | gene-Rv0544c | |
| gORF_106292 | gene-Rv0545c | pitA | gene-Rv0544c | |
| gORF_19021 | gene-Rv0545c | pitA | gene-Rv0543c | |
| gORF_19021 | gene-Rv0545c | pitA | gene-Rv0543c | |
| gORF_19087 | gene-Rv0547c | | gene-Rv0546c | |
| gORF_19087 | gene-Rv0547c | | gene-Rv0546c | |
| gORF_105155 | gene-Rv0584 | | | |
| gORF_105155 | gene-Rv0584 | | | |
| gORF_20570 | gene-Rv0594 | mce2F | gene-Rv0593 | lprL |
| gORF_20570 | gene-Rv0594 | mce2F | gene-Rv0593 | lprL |
| gORF_104161 | gene-Rv0623 | vapB30 | gene-Rv0622 | |
| gORF_104161 | gene-Rv0623 | vapB30 | gene-Rv0622 | |
| gORF_104161 | gene-Rv0624 | vapC30 | | |
| gORF_104161 | gene-Rv0624 | vapC30 | | |
| gORF_103248 | gene-Rv0662c | vapB7 | gene-Rv0661c | vapC7 |
| gORF_103248 | gene-Rv0662c | vapB7 | gene-Rv0661c | vapC7 |
| gORF_23033 | gene-Rv0685 | tuf | | |
| gORF_23033 | gene-Rv0685 | tuf | | |

| | | | | |
|---|---|---|---|---|
| gORF_101684 | gene-Rv0707 | rpsC | | |
| gORF_101684 | gene-Rv0707 | rpsC | | |
| gORF_100972 | gene-Rv0735 | sigL | gene-Rv0734 | mapA |
| gORF_100972 | gene-Rv0735 | sigL | gene-Rv0734 | mapA |
| gORF_26055 | gene-Rv0792c | | | |
| gORF_26055 | gene-Rv0792c | | | |
| gORF_26055 | gene-Rv0793 | | | |
| gORF_26055 | gene-Rv0793 | | | |
| gORF_99466 | gene-Rv0794c | | | |
| gORF_99466 | gene-Rv0794c | | | |
| gORF_26718 | gene-Rv0819 | mshD | | |
| gORF_26718 | gene-Rv0819 | mshD | | |
| gORF_27483 | gene-Rv0841 | | gene-Rv0840c | pip |
| gORF_27483 | gene-Rv0841 | | gene-Rv0840c | pip |
| gORF_98112 | gene-Rv0843 | | gene-Rv0842 | |
| gORF_98112 | gene-Rv0843 | | gene-Rv0842 | |
| gORF_28867 | gene-Rv0887c | | gene-Rv0886 | fprB |
| gORF_28867 | gene-Rv0887c | | gene-Rv0886 | fprB |
| gORF_29058 | gene-Rv0890c | | | |
| gORF_29058 | gene-Rv0890c | | | |
| gORF_29612 | gene-Rv0907 | | gene-Rv0906 | |
| gORF_29612 | gene-Rv0907 | | gene-Rv0906 | |
| gORF_30484 | gene-Rv0931c | pknD | gene-Rv0930 | pstA1 |
| gORF_30484 | gene-Rv0931c | pknD | gene-Rv0930 | pstA1 |
| gORF_31663 | gene-Rv0967 | csoR | gene-Rv0966c | |
| gORF_31663 | gene-Rv0967 | csoR | gene-Rv0966c | |
| gORF_94112 | gene-Rv0973c | accA2 | | |
| gORF_94112 | gene-Rv0973c | accA2 | | |
| gORF_32165 | gene-Rv0980c | PE_PGRS18 | gene-Rv0979c | |
| gORF_32165 | gene-Rv0980c | PE_PGRS18 | gene-Rv0979c | |
| gORF_32272 | gene-Rv0982 | mprB | | |
| gORF_32272 | gene-Rv0982 | mprB | | |
| gORF_92693 | gene-Rv1021 | | gene-Rv1020 | mfd |
| gORF_92693 | gene-Rv1021 | | gene-Rv1020 | mfd |
| gORF_34041 | gene-Rv1040c | PE8 | | |
| gORF_34041 | gene-Rv1040c | PE8 | | |
| gORF_35346 | gene-Rv1078 | | | |
| gORF_35346 | gene-Rv1078 | | | |
| gORF_91014 | gene-Rv1079 | metB | gene-Rv1078 | |
| gORF_91014 | gene-Rv1079 | metB | gene-Rv1078 | |
| gORF_35757 | gene-Rv1091 | PE_PGRS22 | | |
| gORF_35757 | gene-Rv1091 | PE_PGRS22 | | |
| gORF_90691 | gene-Rv1091 | PE_PGRS22 | | |
| gORF_90691 | gene-Rv1091 | PE_PGRS22 | | |
| gORF_35848 | gene-Rv1093 | glyA1 | | |
| gORF_35848 | gene-Rv1093 | glyA1 | | |
| gORF_36517 | gene-Rv1118c | | | |
| gORF_36517 | gene-Rv1118c | | | |
| gORF_36517 | gene-Rv1119c | | | |

| | | | | |
|---|---|---|---|---|
| gORF_36517 | gene-Rv1119c | | | |
| gORF_89122 | gene-Rv1151c | | | |
| gORF_89122 | gene-Rv1151c | | | |
| gORF_86692 | gene-Rv1223 | htrA | | |
| gORF_86692 | gene-Rv1223 | htrA | | |
| gORF_86692 | gene-Rv1224 | tatB | | |
| gORF_86692 | gene-Rv1224 | tatB | | |
| gORF_85045 | gene-Rv1285 | cysD | | |
| gORF_85045 | gene-Rv1285 | cysD | | |
| gORF_42930 | gene-Rv1302 | rfe | gene-Rv1301 | |
| gORF_42930 | gene-Rv1302 | rfe | gene-Rv1301 | |
| gORF_84182 | gene-Rv1308 | atpA | | |
| gORF_84182 | gene-Rv1308 | atpA | | |
| gORF_43305 | gene-Rvnr01 | rrs | gene-Rv1315 | murA |
| gORF_43305 | gene-Rvnr01 | rrs | gene-Rv1315 | murA |
| gORF_83446 | gene-Rv1327c | glgE | | |
| gORF_83446 | gene-Rv1327c | glgE | | |
| gORF_44038 | gene-Rv1327c | glgE | | |
| gORF_44038 | gene-Rv1327c | glgE | | |
| gORF_44524 | gene-Rv1345 | mbtM | | |
| gORF_44524 | gene-Rv1345 | mbtM | | |
| gORF_44871 | gene-Rv1354c | | | |
| gORF_44871 | gene-Rv1354c | | | |
| gORF_44871 | gene-Rv1355c | moeY | | |
| gORF_44871 | gene-Rv1355c | moeY | | |
| gORF_82083 | gene-Rv1366 | | | |
| gORF_82083 | gene-Rv1366 | | | |
| gORF_82083 | gene-Rv1366A | | | |
| gORF_82083 | gene-Rv1366A | | | |
| gORF_81334 | gene-Rv1388 | mihF | gene-Rv1387 | PPE20 |
| gORF_81334 | gene-Rv1388 | mihF | gene-Rv1387 | PPE20 |
| gORF_46630 | gene-Rv1406 | fmt | gene-Rv1405c | |
| gORF_46630 | gene-Rv1406 | fmt | gene-Rv1405c | |
| gORF_80243 | gene-Rv1430 | PE16 | | |
| gORF_80243 | gene-Rv1430 | PE16 | | |
| gORF_47860 | gene-Rv1448c | tal | gene-Rv1447c | zwf2 |
| gORF_47860 | gene-Rv1448c | tal | gene-Rv1447c | zwf2 |
| gORF_48161 | gene-Rv1455 | | gene-Rv1454c | qor |
| gORF_48161 | gene-Rv1455 | | gene-Rv1454c | qor |
| gORF_48466 | gene-Rv1464 | csd | gene-Rv1463 | |
| gORF_48466 | gene-Rv1464 | csd | gene-Rv1463 | |
| gORF_78930 | gene-Rv1468c | PE_PGRS29 | gene-Rv1467c | fadE15 |
| gORF_78930 | gene-Rv1468c | PE_PGRS29 | gene-Rv1467c | fadE15 |
| gORF_78109 | gene-Rv1492 | mutA | gene-Rv1491c | |
| gORF_78109 | gene-Rv1492 | mutA | gene-Rv1491c | |
| gORF_50161 | gene-Rv1515c | | gene-Rv1514c | |
| gORF_50161 | gene-Rv1515c | | gene-Rv1514c | |
| gORF_50213 | gene-Rv1516c | | | |
| gORF_50213 | gene-Rv1516c | | | |

| | | | | |
|---|---|---|---|---|
| gORF_51571 | gene-Rv1548c | PPE21 | | |
| gORF_51571 | gene-Rv1548c | PPE21 | | |
| gORF_75976 | gene-Rv1551 | plsB1 | | |
| gORF_75976 | gene-Rv1551 | plsB1 | | |
| gORF_75871 | gene-Rv1553 | frdB | gene-Rv1552 | frdA |
| gORF_75871 | gene-Rv1553 | frdB | gene-Rv1552 | frdA |
| gORF_52560 | gene-Rv1582c | | | |
| gORF_52560 | gene-Rv1582c | | | |
| gORF_52560 | gene-Rv1583c | | | |
| gORF_52560 | gene-Rv1583c | | | |
| gORF_74745 | gene-Rv1603 | hisA | | |
| gORF_74745 | gene-Rv1603 | hisA | | |
| gORF_74745 | gene-Rv1604 | impA | | |
| gORF_74745 | gene-Rv1604 | impA | | |
| gORF_74533 | gene-Rv1612 | trpB | | |
| gORF_74533 | gene-Rv1612 | trpB | | |
| gORF_53666 | gene-Rv1622c | cydB | gene-Rv1621c | cydD |
| gORF_53666 | gene-Rv1622c | cydB | gene-Rv1621c | cydD |
| gORF_55538 | gene-Rv1672c | | | |
| gORF_55538 | gene-Rv1672c | | | |
| gORF_56439 | gene-Rv1703c | | gene-Rv1702c | |
| gORF_56439 | gene-Rv1703c | | gene-Rv1702c | |
| gORF_56876 | gene-Rv1718 | | gene-Rv1717 | |
| gORF_56876 | gene-Rv1718 | | gene-Rv1717 | |
| gORF_56923 | gene-Rv1720c | vapC12 | gene-Rvnt21 | proT |
| gORF_56923 | gene-Rv1720c | vapC12 | gene-Rvnt21 | proT |
| gORF_69072 | gene-Rv1775 | | | |
| gORF_69072 | gene-Rv1775 | | | |
| gORF_69072 | gene-Rv1776c | | | |
| gORF_69072 | gene-Rv1776c | | | |
| gORF_69007 | gene-Rv1777 | cyp144 | | |
| gORF_69007 | gene-Rv1777 | cyp144 | | |
| gORF_68832 | gene-Rv1782 | eccB5 | | |
| gORF_68832 | gene-Rv1782 | eccB5 | | |
| gORF_68833 | gene-Rv1782 | eccB5 | | |
| gORF_68833 | gene-Rv1782 | eccB5 | | |
| gORF_68829 | gene-Rv1782 | eccB5 | | |
| gORF_68829 | gene-Rv1782 | eccB5 | | |
| gORF_59194 | gene-Rv1785c | cyp143 | | |
| gORF_59194 | gene-Rv1785c | cyp143 | | |
| gORF_68366 | gene-Rv1795 | eccD5 | | |
| gORF_68366 | gene-Rv1795 | eccD5 | | |
| gORF_68366 | gene-Rv1796 | mycP5 | | |
| gORF_68366 | gene-Rv1796 | mycP5 | | |
| gORF_67977 | gene-Rv1802 | PPE30 | | |
| gORF_67977 | gene-Rv1802 | PPE30 | | |
| gORF_67918 | gene-Rv1803c | PE_PGRS32 | | |
| gORF_67918 | gene-Rv1803c | PE_PGRS32 | | |
| gORF_67446 | gene-Rv1817 | | | |

| | | | | |
|---|---|---|---|---|
| gORF_67446 | gene-Rv1817 | | | |
| gORF_67145 | gene-Rv1823 | | gene-Rv1822 | pgsA2 |
| gORF_67145 | gene-Rv1823 | | gene-Rv1822 | pgsA2 |
| gORF_61194 | gene-Rv1844c | gnd1 | | |
| gORF_61194 | gene-Rv1844c | gnd1 | | |
| gORF_65795 | gene-Rv1868 | | | |
| gORF_65795 | gene-Rv1868 | | | |
| gORF_65479 | gene-Rv1877 | | | |
| gORF_65479 | gene-Rv1877 | | | |
| gORF_65479 | gene-Rv1878 | glnA3 | | |
| gORF_65479 | gene-Rv1878 | glnA3 | | |
| gORF_64858 | gene-Rv1901 | cinA | | |
| gORF_64858 | gene-Rv1901 | cinA | | |
| gORF_62933 | gene-Rv1907c | | | |
| gORF_62933 | gene-Rv1907c | | | |
| gORF_63052 | gene-Rv1911c | lppC | gene-Rv1910c | |
| gORF_63052 | gene-Rv1911c | lppC | gene-Rv1910c | |
| gORF_62770 | gene-Rv1974 | | gene-Rv1973 | |
| gORF_62770 | gene-Rv1974 | | gene-Rv1973 | |
| gORF_62770 | gene-Rv1975 | | | |
| gORF_62770 | gene-Rv1975 | | | |
| gORF_64903 | gene-Rv1979c | | | |
| gORF_64903 | gene-Rv1979c | | | |
| gORF_61964 | gene-Rv1997 | ctpF | | |
| gORF_61964 | gene-Rv1997 | ctpF | | |
| gORF_61558 | gene-Rv2006 | otsB1 | | |
| gORF_61558 | gene-Rv2006 | otsB1 | | |
| gORF_66553 | gene-Rv2030c | | | |
| gORF_66553 | gene-Rv2030c | | | |
| gORF_67285 | gene-Rv2048c | pks12 | | |
| gORF_67285 | gene-Rv2048c | pks12 | | |
| gORF_59932 | gene-Rv2052c | | | |
| gORF_59932 | gene-Rv2052c | | | |
| gORF_68412 | gene-Rv2091c | | gene-Rv2090 | |
| gORF_68412 | gene-Rv2091c | | gene-Rv2090 | |
| gORF_68418 | gene-Rv2092c | helY | gene-Rv2091c | |
| gORF_68418 | gene-Rv2092c | helY | gene-Rv2091c | |
| gORF_68556 | gene-Rv2095c | pafC | | |
| gORF_68556 | gene-Rv2095c | pafC | | |
| gORF_57646 | gene-Rv2127 | ansP1 | | |
| gORF_57646 | gene-Rv2127 | ansP1 | | |
| gORF_57646 | gene-Rv2128 | | | |
| gORF_57646 | gene-Rv2128 | | | |
| gORF_57419 | gene-Rv2138 | lppL | gene-Rv2137c | |
| gORF_57419 | gene-Rv2138 | lppL | gene-Rv2137c | |
| gORF_70944 | gene-Rv2182c | | | |
| gORF_70944 | gene-Rv2182c | | | |
| gORF_70944 | gene-Rv2183c | | | |
| gORF_70944 | gene-Rv2183c | | | |

| | | | | |
|---|---|---|---|---|
| gORF_55260 | gene-Rv2212 | | gene-Rv2211c | gcvT |
| gORF_55260 | gene-Rv2212 | | gene-Rv2211c | gcvT |
| gORF_72140 | gene-Rv2223c | | | |
| gORF_72140 | gene-Rv2223c | | | |
| gORF_54773 | gene-Rv2224c | caeA | | |
| gORF_54773 | gene-Rv2224c | caeA | | |
| gORF_72705 | gene-Rv2242 | | | |
| gORF_72705 | gene-Rv2242 | | | |
| gORF_53339 | gene-Rv2268c | cyp128 | gene-Rv2267c | |
| gORF_53339 | gene-Rv2268c | cyp128 | gene-Rv2267c | |
| gORF_52583 | gene-Rv2289 | cdh | gene-Rv2288 | |
| gORF_52583 | gene-Rv2289 | cdh | gene-Rv2288 | |
| gORF_52566 | gene-Rv2290 | lppO | gene-Rv2289 | cdh |
| gORF_52566 | gene-Rv2290 | lppO | gene-Rv2289 | cdh |
| gORF_52031 | gene-Rv2307B | | gene-Rv2307A | |
| gORF_52031 | gene-Rv2307B | | gene-Rv2307A | |
| gORF_51888 | gene-Rv2310 | | gene-Rv2309A | |
| gORF_51888 | gene-Rv2310 | | gene-Rv2309A | |
| gORF_74686 | gene-Rv2319c | | gene-Rv2318 | uspC |
| gORF_74686 | gene-Rv2319c | | gene-Rv2318 | uspC |
| gORF_75042 | gene-Rv2329c | narK1 | | |
| gORF_75042 | gene-Rv2329c | narK1 | | |
| gORF_51269 | gene-Rv2332 | mez | | |
| gORF_51269 | gene-Rv2332 | mez | | |
| gORF_75428 | gene-Rv2339 | mmpL9 | | |
| gORF_75428 | gene-Rv2339 | mmpL9 | | |
| gORF_75773 | gene-Rv2349c | plcC | | |
| gORF_75773 | gene-Rv2349c | plcC | | |
| gORF_75773 | gene-Rv2350c | plcB | | |
| gORF_75773 | gene-Rv2350c | plcB | | |
| gORF_76198 | gene-Rv2357c | glyS | | |
| gORF_76198 | gene-Rv2357c | glyS | | |
| gORF_50133 | gene-Rv2357c | glyS | | |
| gORF_50133 | gene-Rv2357c | glyS | | |
| gORF_48905 | gene-Rv2394 | ggtB | | |
| gORF_48905 | gene-Rv2394 | ggtB | | |
| gORF_48053 | gene-Rv2421c | nadD | gene-Rv2420c | |
| gORF_48053 | gene-Rv2421c | nadD | gene-Rv2420c | |
| gORF_78990 | gene-Rv2441c | rpmA | gene-Rv2440c | obg |
| gORF_78990 | gene-Rv2441c | rpmA | gene-Rv2440c | obg |
| gORF_78990 | gene-Rv2442c | rplU | | |
| gORF_78990 | gene-Rv2442c | rplU | | |
| gORF_46723 | gene-Rv2466c | | | |
| gORF_46723 | gene-Rv2466c | | | |
| gORF_46723 | gene-Rv2467 | pepN | | |
| gORF_46723 | gene-Rv2467 | pepN | | |
| gORF_80264 | gene-Rv2478c | | gene-Rv2477c | |
| gORF_80264 | gene-Rv2478c | | gene-Rv2477c | |
| gORF_46319 | gene-Rv2478c | | gene-Rv2477c | |

| | | | | |
|---|---|---|---|---|
| gORF_46319 | gene-Rv2478c | | gene-Rv2477c | |
| gORF_45994 | gene-Rv2488c | | | |
| gORF_45994 | gene-Rv2488c | | | |
| gORF_45925 | gene-Rv2490c | PE_PGRS43 | gene-Rv2489c | |
| gORF_45925 | gene-Rv2490c | PE_PGRS43 | gene-Rv2489c | |
| gORF_45808 | gene-Rv2490c | PE_PGRS43 | | |
| gORF_45808 | gene-Rv2490c | PE_PGRS43 | | |
| gORF_83262 | gene-Rv2568c | | | |
| gORF_83262 | gene-Rv2568c | | | |
| gORF_43180 | gene-Rv2579 | dhaA | gene-Rv2578c | |
| gORF_43180 | gene-Rv2579 | dhaA | gene-Rv2578c | |
| gORF_42594 | gene-Rv2598 | | gene-Rv2597 | |
| gORF_42594 | gene-Rv2598 | | gene-Rv2597 | |
| gORF_42379 | gene-Rv2606c | snzP | gene-Rv2605c | tesB2 |
| gORF_42379 | gene-Rv2606c | snzP | gene-Rv2605c | tesB2 |
| gORF_42096 | gene-Rv2614A | | gene-Rv2614c | thrS |
| gORF_42096 | gene-Rv2614A | | gene-Rv2614c | thrS |
| gORF_42096 | gene-Rv2615c | PE_PGRS45 | | |
| gORF_42096 | gene-Rv2615c | PE_PGRS45 | | |
| gORF_84771 | gene-Rv2624c | | | |
| gORF_84771 | gene-Rv2624c | | | |
| gORF_84771 | gene-Rv2625c | | | |
| gORF_84771 | gene-Rv2625c | | | |
| gORF_84877 | gene-Rv2627c | | | |
| gORF_84877 | gene-Rv2627c | | | |
| gORF_85060 | gene-Rv2632c | | gene-Rv2631 | |
| gORF_85060 | gene-Rv2632c | | gene-Rv2631 | |
| gORF_41235 | gene-Rvnt31 | valT | | |
| gORF_41235 | gene-Rvnt31 | valT | | |
| gORF_85603 | gene-Rv2659c | | | |
| gORF_85603 | gene-Rv2659c | | | |
| gORF_40039 | gene-Rv2693c | | gene-Rv2692 | ceoC |
| gORF_40039 | gene-Rv2693c | | gene-Rv2692 | ceoC |
| gORF_38811 | gene-Rv2737c | recA | | |
| gORF_38811 | gene-Rv2737c | recA | | |
| gORF_38583 | gene-Rv2745c | clgR | gene-Rv2744c | 35kd_ag |
| gORF_38583 | gene-Rv2745c | clgR | gene-Rv2744c | 35kd_ag |
| gORF_87581 | gene-Rv2747 | argA | gene-Rv2746c | pgsA3 |
| gORF_87581 | gene-Rv2747 | argA | gene-Rv2746c | pgsA3 |
| gORF_87844 | gene-Rv2756c | hsdM | | |
| gORF_87844 | gene-Rv2756c | hsdM | | |
| gORF_87937 | gene-Rv2761c | hsdS | | |
| gORF_87937 | gene-Rv2761c | hsdS | | |
| gORF_38000 | gene-Rv2771c | | | |
| gORF_38000 | gene-Rv2771c | | | |
| gORF_38000 | gene-Rv2772c | | | |
| gORF_38000 | gene-Rv2772c | | | |
| gORF_88422 | gene-Rv2781c | | | |
| gORF_88422 | gene-Rv2781c | | | |

| | | | | |
|---|---|---|---|---|
| gORF_37048 | gene-Rv2812 | | | |
| gORF_37048 | gene-Rv2812 | | | |
| gORF_35817 | gene-Rv2857c | | | |
| gORF_35817 | gene-Rv2857c | | | |
| gORF_90956 | gene-Rv2862c | | gene-Rv2861c | mapB |
| gORF_90956 | gene-Rv2862c | | gene-Rv2861c | mapB |
| gORF_35414 | gene-Rv2873 | mpt83 | gene-Rv2872 | vapC43 |
| gORF_35414 | gene-Rv2873 | mpt83 | gene-Rv2872 | vapC43 |
| gORF_91428 | gene-Rv2881c | cdsA | | |
| gORF_91428 | gene-Rv2881c | cdsA | | |
| gORF_92020 | gene-Rv2902c | rnhB | gene-Rv2901c | |
| gORF_92020 | gene-Rv2902c | rnhB | gene-Rv2901c | |
| gORF_93150 | gene-Rv2933 | ppsC | | |
| gORF_93150 | gene-Rv2933 | ppsC | | |
| gORF_33042 | gene-Rv2934 | ppsD | | |
| gORF_33042 | gene-Rv2934 | ppsD | | |
| gORF_32818 | gene-Rv2935 | ppsE | | |
| gORF_32818 | gene-Rv2935 | ppsE | | |
| gORF_31439 | gene-Rv2961 | | gene-Rv2960c | |
| gORF_31439 | gene-Rv2961 | | gene-Rv2960c | |
| gORF_95002 | gene-Rv2975a | | gene-Rv2974c | |
| gORF_95002 | gene-Rv2975a | | gene-Rv2974c | |
| gORF_95424 | gene-Rv2990c | | | |
| gORF_95424 | gene-Rv2990c | | | |
| gORF_95760 | gene-Rv3000 | | gene-Rv2999 | lppY |
| gORF_95760 | gene-Rv3000 | | gene-Rv2999 | lppY |
| gORF_29356 | gene-Rv3025c | iscS | | |
| gORF_29356 | gene-Rv3025c | iscS | | |
| gORF_28422 | gene-Rv3059 | cyp136 | gene-Rv3058c | |
| gORF_28422 | gene-Rv3059 | cyp136 | gene-Rv3058c | |
| gORF_27035 | gene-Rv3096 | | | |
| gORF_27035 | gene-Rv3096 | | | |
| gORF_26752 | gene-Rv3107c | agpS | | |
| gORF_26752 | gene-Rv3107c | agpS | | |
| gORF_26725 | gene-Rv3108 | | gene-Rv3107c | agpS |
| gORF_26725 | gene-Rv3108 | | gene-Rv3107c | agpS |
| gORF_25839 | gene-Rv3136 | PPE51 | | |
| gORF_25839 | gene-Rv3136 | PPE51 | | |
| gORF_25784 | gene-Rv3137 | | | |
| gORF_25784 | gene-Rv3137 | | | |
| gORF_25784 | gene-Rv3138 | pflA | | |
| gORF_25784 | gene-Rv3138 | pflA | | |
| gORF_99913 | gene-Rv3144c | PPE52 | gene-Rv3143 | |
| gORF_99913 | gene-Rv3144c | PPE52 | gene-Rv3143 | |
| gORF_101064 | gene-Rv3190c | | | |
| gORF_101064 | gene-Rv3190c | | | |
| gORF_101587 | gene-Rv3201c | | | |
| gORF_101587 | gene-Rv3201c | | | |
| gORF_102158 | gene-Rv3221A | rshA | gene-Rv3221c | TB7.3 |

| | | | | |
|---|---|---|---|---|
| gORF_102158 | gene-Rv3221A | rshA | gene-Rv3221c | TB7.3 |
| gORF_102158 | gene-Rv3222c | | | |
| gORF_102158 | gene-Rv3222c | | | |
| gORF_102652 | gene-Rv3239c | | | |
| gORF_102652 | gene-Rv3239c | | | |
| gORF_103423 | gene-Rv3263 | | | |
| gORF_103423 | gene-Rv3263 | | | |
| gORF_103642 | gene-Rv3271c | | gene-Rv3270 | ctpC |
| gORF_103642 | gene-Rv3271c | | gene-Rv3270 | ctpC |
| gORF_20316 | gene-Rv3307 | deoD | | |
| gORF_20316 | gene-Rv3307 | deoD | | |
| gORF_20316 | gene-Rv3308 | pmmB | | |
| gORF_20316 | gene-Rv3308 | pmmB | | |
| gORF_104937 | gene-Rv3322c | | gene-Rv3320c | vapC44 |
| gORF_104937 | gene-Rv3322c | | gene-Rv3320c | vapC44 |
| gORF_19681 | gene-Rv3328c | sigJ | | |
| gORF_19681 | gene-Rv3328c | sigJ | | |
| gORF_19557 | gene-Rv3332 | nagA | gene-Rv3331 | sugI |
| gORF_19557 | gene-Rv3332 | nagA | gene-Rv3331 | sugI |
| gORF_19418 | gene-Rv3336c | trpS | | |
| gORF_19418 | gene-Rv3336c | trpS | | |
| gORF_105261 | gene-Rv3336c | trpS | gene-Rv3335c | |
| gORF_105261 | gene-Rv3336c | trpS | gene-Rv3335c | |
| gORF_105604 | gene-Rv3343c | PPE54 | | |
| gORF_105604 | gene-Rv3343c | PPE54 | | |
| gORF_105783 | gene-Rv3343c | PPE54 | | |
| gORF_105783 | gene-Rv3343c | PPE54 | | |
| gORF_105910 | gene-Rv3346c | | gene-Rv3345c | PE_PGRS50 |
| gORF_105910 | gene-Rv3346c | | gene-Rv3345c | PE_PGRS50 |
| gORF_106326 | gene-Rv3347c | PPE55 | | |
| gORF_106326 | gene-Rv3347c | PPE55 | | |
| gORF_106880 | gene-Rv3350c | PPE56 | | |
| gORF_106880 | gene-Rv3350c | PPE56 | | |
| gORF_106973 | gene-Rv3350c | PPE56 | | |
| gORF_106973 | gene-Rv3350c | PPE56 | | |
| gORF_107229 | gene-Rv3365c | | | |
| gORF_107229 | gene-Rv3365c | | | |
| gORF_107491 | gene-Rv3371 | | | |
| gORF_107491 | gene-Rv3371 | | | |
| gORF_107491 | gene-Rv3372 | otsB2 | | |
| gORF_107491 | gene-Rv3372 | otsB2 | | |
| gORF_109217 | gene-Rv3429 | PPE59 | | |
| gORF_109217 | gene-Rv3429 | PPE59 | | |
| gORF_109217 | gene-Rv3430c | | | |
| gORF_109217 | gene-Rv3430c | | | |
| gORF_16021 | gene-Rv3437 | | gene-Rv3436c | glmS |
| gORF_16021 | gene-Rv3437 | | gene-Rv3436c | glmS |
| gORF_14677 | gene-Rv3483c | | | |
| gORF_14677 | gene-Rv3483c | | | |

| | | | | |
|---|---|---|---|---|
| gORF_14611 | gene-Rv3484 | | | |
| gORF_14611 | gene-Rv3484 | | | |
| gORF_14254 | gene-Rv3500c | yrbE4B | gene-Rv3499c | mce4A |
| gORF_14254 | gene-Rv3500c | yrbE4B | gene-Rv3499c | mce4A |
| gORF_111749 | gene-Rv3514 | PE_PGRS57 | | |
| gORF_111749 | gene-Rv3514 | PE_PGRS57 | | |
| gORF_112725 | gene-Rv3544c | fadE28 | | |
| gORF_112725 | gene-Rv3544c | fadE28 | | |
| gORF_11817 | gene-Rv3576 | lppH | | |
| gORF_11817 | gene-Rv3576 | lppH | | |
| gORF_113929 | gene-Rv3590c | PE_PGRS58 | | |
| gORF_113929 | gene-Rv3590c | PE_PGRS58 | | |
| gORF_114887 | gene-Rv3627c | | | |
| gORF_114887 | gene-Rv3627c | | | |
| gORF_9109 | gene-Rv3670 | ephE | | |
| gORF_9109 | gene-Rv3670 | ephE | | |
| gORF_7579 | gene-Rv3717 | | gene-Rv3716c | |
| gORF_7579 | gene-Rv3717 | | gene-Rv3716c | |
| gORF_7258 | gene-Rv3726 | | | |
| gORF_7258 | gene-Rv3726 | | | |
| gORF_117748 | gene-Rv3729 | | | |
| gORF_117748 | gene-Rv3729 | | | |
| gORF_6190 | gene-Rv3759c | proX | gene-Rv3758c | proV |
| gORF_6190 | gene-Rv3759c | proX | gene-Rv3758c | proV |
| gORF_118806 | gene-Rv3770c | | | |
| gORF_118806 | gene-Rv3770c | | | |
| gORF_5683 | gene-Rv3776 | | | |
| gORF_5683 | gene-Rv3776 | | | |
| gORF_119727 | gene-Rv3801c | fadD32 | | |
| gORF_119727 | gene-Rv3801c | fadD32 | | |
| gORF_120720 | gene-Rv3825c | pks2 | | |
| gORF_120720 | gene-Rv3825c | pks2 | | |
| gORF_2540 | gene-Rv3852 | hns | gene-Rv3850 | |
| gORF_2540 | gene-Rv3852 | hns | gene-Rv3850 | |
| gORF_2129 | gene-Rv3864 | espE | gene-Rv3863 | |
| gORF_2129 | gene-Rv3864 | espE | gene-Rv3863 | |
| gORF_2002 | gene-Rv3868 | eccA1 | | |
| gORF_2002 | gene-Rv3868 | eccA1 | | |
| gORF_1756 | gene-Rv3871 | eccCb1 | | |
| gORF_1756 | gene-Rv3871 | eccCb1 | | |
| gORF_122227 | gene-Rv3876 | espI | | |
| gORF_122227 | gene-Rv3876 | espI | | |
| gORF_443 | gene-Rv3910 | | | |
| gORF_443 | gene-Rv3910 | | | |
| gORF_376 | gene-Rv3910 | | | |
| gORF_376 | gene-Rv3910 | | | |
| gORF_329 | gene-Rv3912 | | gene-Rv3911 | sigM |
| gORF_329 | gene-Rv3912 | | gene-Rv3911 | sigM |
| tORF_288 | gene-Rv0006 | gyrA | | |

| | | | | |
|---|---|---|---|---|
| tORF_288 | gene-Rv0006 | gyrA | | |
| tORF_438 | gene-Rv0014c | pknB | | |
| tORF_438 | gene-Rv0014c | pknB | | |
| tORF_437 | gene-Rv0014c | pknB | | |
| tORF_437 | gene-Rv0014c | pknB | | |
| tORF_433 | gene-Rv0014c | pknB | | |
| tORF_433 | gene-Rv0014c | pknB | | |
| tORF_421 | gene-Rv0014c | pknB | gene-Rv0013 | trpG |
| tORF_421 | gene-Rv0014c | pknB | gene-Rv0013 | trpG |
| tORF_2047 | gene-Rv0069c | sdaA | | |
| tORF_2047 | gene-Rv0069c | sdaA | | |
| tORF_3074 | gene-Rv0101 | nrp | | |
| tORF_3074 | gene-Rv0101 | nrp | | |
| tORF_3622 | gene-Rv0115a | | gene-Rv0115 | hddA |
| tORF_3622 | gene-Rv0115a | | gene-Rv0115 | hddA |
| tORF_3808 | gene-Rv0121c | | gene-Rv0120c | fusA2 |
| tORF_3808 | gene-Rv0121c | | gene-Rv0120c | fusA2 |
| tORF_3869 | gene-Rv0124 | PE_PGRS2 | | |
| tORF_3869 | gene-Rv0124 | PE_PGRS2 | | |
| tORF_4175 | gene-Rv0139 | | | |
| tORF_4175 | gene-Rv0139 | | | |
| tORF_4671 | gene-Rv0163 | | gene-Rv0162c | adhE1 |
| tORF_4671 | gene-Rv0163 | | gene-Rv0162c | adhE1 |
| tORF_7075 | gene-Rv0212c | nadR | | |
| tORF_7075 | gene-Rv0212c | nadR | | |
| tORF_7075 | gene-Rv0213c | | | |
| tORF_7075 | gene-Rv0213c | | | |
| tORF_8639 | gene-Rv0278c | | gene-Rv0277c | vapC25 |
| tORF_8639 | gene-Rv0278c | | gene-Rv0277c | vapC25 |
| tORF_8883 | gene-Rv0284 | eccC3 | | |
| tORF_8883 | gene-Rv0284 | eccC3 | | |
| tORF_9636 | gene-Rv0309 | | | |
| tORF_9636 | gene-Rv0309 | | | |
| tORF_9734 | gene-Rv0315 | | gene-Rv0314c | |
| tORF_9734 | gene-Rv0315 | | gene-Rv0314c | |
| tORF_10760 | gene-Rv0355c | PPE8 | | |
| tORF_10760 | gene-Rv0355c | PPE8 | | |
| tORF_11163 | gene-Rv0371c | | gene-Rv0370c | |
| tORF_11163 | gene-Rv0371c | | gene-Rv0370c | |
| tORF_11242 | gene-Rv0373c | | gene-Rv0372c | |
| tORF_11242 | gene-Rv0373c | | gene-Rv0372c | |
| tORF_11233 | gene-Rv0373c | | | |
| tORF_11233 | gene-Rv0373c | | | |
| tORF_12232 | gene-Rv0405 | pks6 | | |
| tORF_12232 | gene-Rv0405 | pks6 | | |
| tORF_12940 | gene-Rv0427c | xthA | | |
| tORF_12940 | gene-Rv0427c | xthA | | |
| tORF_14178 | gene-Rv0484c | | gene-Rv0483 | lprQ |
| tORF_14178 | gene-Rv0484c | | gene-Rv0483 | lprQ |

| | | | | |
|---|---|---|---|---|
| tORF_16103 | gene-Rv0545c | pitA | gene-Rv0544c | |
| tORF_16103 | gene-Rv0545c | pitA | gene-Rv0544c | |
| tORF_17297 | gene-Rv0594 | mce2F | gene-Rv0593 | lprL |
| tORF_17297 | gene-Rv0594 | mce2F | gene-Rv0593 | lprL |
| tORF_19112 | gene-Rv0685 | tuf | | |
| tORF_19112 | gene-Rv0685 | tuf | | |
| tORF_20207 | gene-Rv0739 | | | |
| tORF_20207 | gene-Rv0739 | | | |
| tORF_20553 | gene-Rv0756c | | | |
| tORF_20553 | gene-Rv0756c | | | |
| tORF_21472 | gene-Rv0794c | | | |
| tORF_21472 | gene-Rv0794c | | | |
| tORF_22109 | gene-Rv0819 | mshD | | |
| tORF_22109 | gene-Rv0819 | mshD | | |
| tORF_22610 | gene-Rv0843 | | gene-Rv0842 | |
| tORF_22610 | gene-Rv0843 | | gene-Rv0842 | |
| tORF_22787 | gene-Rv0845 | | gene-Rv0844c | narL |
| tORF_22787 | gene-Rv0845 | | gene-Rv0844c | narL |
| tORF_24534 | gene-Rv0907 | | gene-Rv0906 | |
| tORF_24534 | gene-Rv0907 | | gene-Rv0906 | |
| tORF_26189 | gene-Rv0967 | csoR | gene-Rv0966c | |
| tORF_26189 | gene-Rv0967 | csoR | gene-Rv0966c | |
| tORF_26369 | gene-Rv0973c | accA2 | | |
| tORF_26369 | gene-Rv0973c | accA2 | | |
| tORF_26465 | gene-Rv0977 | PE_PGRS16 | | |
| tORF_26465 | gene-Rv0977 | PE_PGRS16 | | |
| tORF_27069 | gene-Rv0997a | | gene-Rv0996 | |
| tORF_27069 | gene-Rv0997a | | gene-Rv0996 | |
| tORF_28759 | gene-Rv1078 | | | |
| tORF_28759 | gene-Rv1078 | | | |
| tORF_29093 | gene-Rv1093 | glyA1 | | |
| tORF_29093 | gene-Rv1093 | glyA1 | | |
| tORF_29123 | gene-Rv1095 | phoH2 | gene-Rv1094 | desA2 |
| tORF_29123 | gene-Rv1095 | phoH2 | gene-Rv1094 | desA2 |
| tORF_29232 | gene-Rv1100 | | gene-Rv1099c | glpX |
| tORF_29232 | gene-Rv1100 | | gene-Rv1099c | glpX |
| tORF_30261 | gene-Rv1151c | | | |
| tORF_30261 | gene-Rv1151c | | | |
| tORF_30438 | gene-Rv1158c | | | |
| tORF_30438 | gene-Rv1158c | | | |
| tORF_30521 | gene-Rv1161 | narG | | |
| tORF_30521 | gene-Rv1161 | narG | | |
| tORF_33916 | gene-Rv1286 | | gene-Rv1285 | cysD |
| tORF_33916 | gene-Rv1286 | | gene-Rv1285 | cysD |
| tORF_33963 | gene-Rv1288 | | gene-Rv1287 | |
| tORF_33963 | gene-Rv1288 | | gene-Rv1287 | |
| tORF_34630 | gene-Rv1315 | murA | | |
| tORF_34630 | gene-Rv1315 | murA | | |
| tORF_35175 | gene-Rv1327c | glgE | | |

| | | | | |
|---|---|---|---|---|
| tORF_35175 | gene-Rv1327c | glgE | | |
| tORF_35369 | gene-Rv1333 | | gene-Rv1332 | |
| tORF_35369 | gene-Rv1333 | | gene-Rv1332 | |
| tORF_35700 | gene-Rv1345 | mbtM | | |
| tORF_35700 | gene-Rv1345 | mbtM | | |
| tORF_36024 | gene-Rv1354c | | | |
| tORF_36024 | gene-Rv1354c | | | |
| tORF_36126 | gene-Rv1355c | moeY | gene-Rv1354c | |
| tORF_36126 | gene-Rv1355c | moeY | gene-Rv1354c | |
| tORF_37787 | gene-Rv1402 | priA | | |
| tORF_37787 | gene-Rv1402 | priA | | |
| tORF_38551 | gene-Rv1428c | | | |
| tORF_38551 | gene-Rv1428c | | | |
| tORF_38722 | gene-Rv1430 | PE16 | | |
| tORF_38722 | gene-Rv1430 | PE16 | | |
| tORF_39731 | gene-Rv1464 | csd | gene-Rv1463 | |
| tORF_39731 | gene-Rv1464 | csd | gene-Rv1463 | |
| tORF_39817 | gene-Rv1468c | PE_PGRS29 | gene-Rv1467c | fadE15 |
| tORF_39817 | gene-Rv1468c | PE_PGRS29 | gene-Rv1467c | fadE15 |
| tORF_39816 | gene-Rv1468c | PE_PGRS29 | gene-Rv1467c | fadE15 |
| tORF_39816 | gene-Rv1468c | PE_PGRS29 | gene-Rv1467c | fadE15 |
| tORF_42121 | gene-Rv1550 | fadD11 | | |
| tORF_42121 | gene-Rv1550 | fadD11 | | |
| tORF_45152 | gene-Rv1657 | argR | gene-Rv1656 | argF |
| tORF_45152 | gene-Rv1657 | argR | gene-Rv1656 | argF |
| tORF_47070 | gene-Rv1718 | | gene-Rv1717 | |
| tORF_47070 | gene-Rv1718 | | gene-Rv1717 | |
| tORF_47766 | gene-Rv1750c | fadD1 | | |
| tORF_47766 | gene-Rv1750c | fadD1 | | |
| tORF_47960 | gene-Rv1759c | wag22 | | |
| tORF_47960 | gene-Rv1759c | wag22 | | |
| tORF_49271 | gene-Rv1803c | PE_PGRS32 | | |
| tORF_49271 | gene-Rv1803c | PE_PGRS32 | | |
| tORF_49343 | gene-Rv1808 | PPE32 | | |
| tORF_49343 | gene-Rv1808 | PPE32 | | |
| tORF_51407 | gene-Rv1899c | lppD | | |
| tORF_51407 | gene-Rv1899c | lppD | | |
| tORF_51407 | gene-Rv1900c | lipJ | | |
| tORF_51407 | gene-Rv1900c | lipJ | | |
| tORF_53372 | gene-Rv1979c | | | |
| tORF_53372 | gene-Rv1979c | | | |
| tORF_53851 | gene-Rv1997 | ctpF | | |
| tORF_53851 | gene-Rv1997 | ctpF | | |
| tORF_55810 | gene-Rv2052c | | | |
| tORF_55810 | gene-Rv2052c | | | |
| tORF_56419 | gene-Rv2075c | | | |
| tORF_56419 | gene-Rv2075c | | | |
| tORF_60096 | gene-Rv2224c | caeA | | |
| tORF_60096 | gene-Rv2224c | caeA | | |

| | | | | |
|---|---|---|---|---|
| tORF_60406 | gene-Rv2234 | ptpA | gene-Rv2232 | ptkA |
| tORF_60406 | gene-Rv2234 | ptpA | gene-Rv2232 | ptkA |
| tORF_61297 | gene-Rv2268c | cyp128 | gene-Rv2267c | |
| tORF_61297 | gene-Rv2268c | cyp128 | gene-Rv2267c | |
| tORF_63427 | gene-Rv2357c | glyS | | |
| tORF_63427 | gene-Rv2357c | glyS | | |
| tORF_65674 | gene-Rv2418c | | gene-Rv2417c | |
| tORF_65674 | gene-Rv2418c | | gene-Rv2417c | |
| tORF_66504 | gene-Rv2444c | rne | | |
| tORF_66504 | gene-Rv2444c | rne | | |
| tORF_67492 | gene-Rv2477c | | | |
| tORF_67492 | gene-Rv2477c | | | |
| tORF_67521 | gene-Rv2478c | | gene-Rv2477c | |
| tORF_67521 | gene-Rv2478c | | gene-Rv2477c | |
| tORF_67823 | gene-Rv2488c | | | |
| tORF_67823 | gene-Rv2488c | | | |
| tORF_67974 | gene-Rv2490c | PE_PGRS43 | gene-Rv2489c | |
| tORF_67974 | gene-Rv2490c | PE_PGRS43 | gene-Rv2489c | |
| tORF_67857 | gene-Rv2490c | PE_PGRS43 | | |
| tORF_67857 | gene-Rv2490c | PE_PGRS43 | | |
| tORF_67941 | gene-Rv2490c | PE_PGRS43 | | |
| tORF_67941 | gene-Rv2490c | PE_PGRS43 | | |
| tORF_68868 | gene-Rv2522c | | gene-Rv2521 | bcp |
| tORF_68868 | gene-Rv2522c | | gene-Rv2521 | bcp |
| tORF_69156 | gene-Rv2530c | vapC39 | gene-Rv2529 | |
| tORF_69156 | gene-Rv2530c | vapC39 | gene-Rv2529 | |
| tORF_69185 | gene-Rv2531c | | | |
| tORF_69185 | gene-Rv2531c | | | |
| tORF_69957 | gene-Rv2566 | | | |
| tORF_69957 | gene-Rv2566 | | | |
| tORF_70947 | gene-Rv2606c | snzP | gene-Rv2605c | tesB2 |
| tORF_70947 | gene-Rv2606c | snzP | gene-Rv2605c | tesB2 |
| tORF_71284 | gene-Rv2615c | PE_PGRS45 | | |
| tORF_71284 | gene-Rv2615c | PE_PGRS45 | | |
| tORF_72141 | gene-Rv2659c | | | |
| tORF_72141 | gene-Rv2659c | | | |
| tORF_72971 | gene-Rv2693c | | gene-Rv2692 | ceoC |
| tORF_72971 | gene-Rv2693c | | gene-Rv2692 | ceoC |
| tORF_73702 | gene-Rv2733c | | gene-Rv2732c | |
| tORF_73702 | gene-Rv2733c | | gene-Rv2732c | |
| tORF_73986 | gene-Rv2737c | recA | | |
| tORF_73986 | gene-Rv2737c | recA | | |
| tORF_74194 | gene-Rv2747 | argA | gene-Rv2746c | pgsA3 |
| tORF_74194 | gene-Rv2747 | argA | gene-Rv2746c | pgsA3 |
| tORF_74872 | gene-Rv2781c | | gene-Rv2780 | ald |
| tORF_74872 | gene-Rv2781c | | gene-Rv2780 | ald |
| tORF_75796 | gene-Rv2826c | | | |
| tORF_75796 | gene-Rv2826c | | | |
| tORF_76226 | gene-Rv2842c | | gene-Rv2841c | nusA |

| | | | | |
|---|---|---|---|---|
| tORF_76226 | gene-Rv2842c | | gene-Rv2841c | nusA |
| tORF_76742 | gene-Rv2857c | | | |
| tORF_76742 | gene-Rv2857c | | | |
| tORF_79117 | gene-Rv2933 | ppsC | | |
| tORF_79117 | gene-Rv2933 | ppsC | | |
| tORF_80148 | gene-Rv2948c | fadD22 | | |
| tORF_80148 | gene-Rv2948c | fadD22 | | |
| tORF_80934 | gene-Rv2975a | | gene-Rv2974c | |
| tORF_80934 | gene-Rv2975a | | gene-Rv2974c | |
| tORF_81913 | gene-Rv3014c | ligA | | |
| tORF_81913 | gene-Rv3014c | ligA | | |
| tORF_82961 | gene-Rv3061c | fadE22 | | |
| tORF_82961 | gene-Rv3061c | fadE22 | | |
| tORF_84082 | gene-Rv3107c | agpS | | |
| tORF_84082 | gene-Rv3107c | agpS | | |
| tORF_84912 | gene-Rv3139 | fadE24 | gene-Rv3138 | pflA |
| tORF_84912 | gene-Rv3139 | fadE24 | gene-Rv3138 | pflA |
| tORF_87534 | gene-Rv3240c | secA1 | | |
| tORF_87534 | gene-Rv3240c | secA1 | | |
| tORF_87698 | gene-Rv3245c | mtrB | gene-Rv3244c | lpqB |
| tORF_87698 | gene-Rv3245c | mtrB | gene-Rv3244c | lpqB |
| tORF_88156 | gene-Rv3263 | | | |
| tORF_88156 | gene-Rv3263 | | | |
| tORF_88201 | gene-Rv3264c | manB | gene-Rv3263 | |
| tORF_88201 | gene-Rv3264c | manB | gene-Rv3263 | |
| tORF_88334 | gene-Rv3271c | | gene-Rv3270 | ctpC |
| tORF_88334 | gene-Rv3271c | | gene-Rv3270 | ctpC |
| tORF_89037 | gene-Rv3293 | pcd | | |
| tORF_89037 | gene-Rv3293 | pcd | | |
| tORF_89298 | gene-Rv3303c | lpdA | | |
| tORF_89298 | gene-Rv3303c | lpdA | | |
| tORF_89943 | gene-Rv3328c | sigJ | | |
| tORF_89943 | gene-Rv3328c | sigJ | | |
| tORF_90137 | gene-Rv3336c | trpS | | |
| tORF_90137 | gene-Rv3336c | trpS | | |
| tORF_90234 | gene-Rv3342 | | gene-Rv3341 | metA |
| tORF_90234 | gene-Rv3342 | | gene-Rv3341 | metA |
| tORF_92272 | gene-Rv3413c | | | |
| tORF_92272 | gene-Rv3413c | | | |
| tORF_92330 | gene-Rv3416 | whiB3 | gene-Rv3415c | |
| tORF_92330 | gene-Rv3416 | whiB3 | gene-Rv3415c | |
| tORF_92637 | gene-Rv3429 | PPE59 | | |
| tORF_92637 | gene-Rv3429 | PPE59 | | |
| tORF_92637 | gene-Rv3430c | | | |
| tORF_92637 | gene-Rv3430c | | | |
| tORF_94315 | gene-Rv3500c | yrbE4B | gene-Rv3499c | mce4A |
| tORF_94315 | gene-Rv3500c | yrbE4B | gene-Rv3499c | mce4A |
| tORF_94745 | gene-Rv3514 | PE_PGRS57 | | |
| tORF_94745 | gene-Rv3514 | PE_PGRS57 | | |

| | | | | |
|---|---|---|---|---|
| tORF_95238 | gene-Rv3537 | kstD | | |
| tORF_95238 | gene-Rv3537 | kstD | | |
| tORF_98881 | gene-Rv3678c | | gene-Rv3677c | |
| tORF_98881 | gene-Rv3678c | | gene-Rv3677c | |
| tORF_99968 | gene-Rv3721c | dnaZX | | |
| tORF_99968 | gene-Rv3721c | dnaZX | | |
| tORF_100857 | gene-Rv3759c | proX | gene-Rv3758c | proV |
| tORF_100857 | gene-Rv3759c | proX | gene-Rv3758c | proV |
| tORF_102211 | gene-Rv3811 | | | |
| tORF_102211 | gene-Rv3811 | | | |
| tORF_102203 | gene-Rv3811 | | | |
| tORF_102203 | gene-Rv3811 | | | |
| tORF_103255 | gene-Rv3848 | | gene-Rv3847 | |
| tORF_103255 | gene-Rv3848 | | gene-Rv3847 | |
| tORF_103888 | gene-Rv3876 | espI | | |
| tORF_103888 | gene-Rv3876 | espI | | |
| tORF_104672 | gene-Rv3897c | | gene-Rv3896c | |
| tORF_104672 | gene-Rv3897c | | gene-Rv3896c | |
| tORF_104870 | gene-Rv3903c | | | |
| tORF_104870 | gene-Rv3903c | | | |

Supplementary Table 3. smORF essentiality. E = Essential, GD = Growth Defect, GA = Growth Advantage

| **smORFs** | **Call** |
|---|---|
| tORF_87698 | GD |
| gORF_250 | ES |
| gORF_15589 | ES |
| gORF_43305 | ES |
| gORF_65479 | GA |
| gORF_16021 | GA |
| gORF_28297 | GA |
| gORF_109100 | ES |
| gORF_70944 | ES |
| gORF_48161 | GA |
| gORF_43305 | ES |
| gORF_105910 | GA |
| gORF_77518 | ES |

Supplementary Table 4. Differentially expressed tORFs upon exposure to antibiotics. Available as a separate Excel file (see Supplementary Materials).

Supplementary Table 5: UpSet groups.

| **group** | **smorfs** |
|---|---|
| isoniazid_24h | tORF_9636 |
| isoniazid_24h | tORF_56419 |
| isoniazid_24h | tORF_60096 |
| isoniazid_24h | tORF_61297 |
| isoniazid_24h | tORF_73986 |
| isoniazid_24h,isoniazid_72h | tORF_20553 |
| isoniazid_24h,isoniazid_72h | tORFs_39817_39816 |
| isoniazid_24h,isoniazid_72h | tORF_76226 |
| isoniazid_24h,isoniazid_72h,pyranizoic_72h | tORF_22610 |
| isoniazid_24h,isoniazid_72h,pyranizoic_04h,pyranizoic_72h,rifampicin_72h | tORF_39731 |
| isoniazid_24h,isoniazid_72h,pyranizoic_24h,pyranizoic_72h,rifampicin_24h,rifampicin_04h | tORF_49343 |
| isoniazid_24h,isoniazid_04h,rifampicin_24h,rifampicin_04h,rifampicin_72h | tORF_53851 |
| isoniazid_24h,isoniazid_72h,rifampicin_04h,rifampicin_72h | tORF_55810 |
| isoniazid_24h,isoniazid_72h,rifampicin_24h,rifampicin_72h | tORF_72971 |
| isoniazid_24h,isoniazid_72h,pyranizoic_72h,rifampicin_72h | tORF_88334 |
| isoniazid_24h,isoniazid_04h | tORF_92272 |
| isoniazid_24h,isoniazid_72h,pyranizoic_24h,pyranizoic_04h,pyranizoic_72h,rifampicin_24h,rifampicin_72h | tORF_92330 |
| isoniazid_24h,isoniazid_72h,pyranizoic_72h,rifampicin_24h,rifampicin_72h | tORF_103255 |
| isoniazid_04h,rifampicin_04h | tORF_30438 |
| isoniazid_72h,rifampicin_04h | tORF_3622 |
| isoniazid_72h | tORF_3869 |
| isoniazid_72h | tORF_8883 |
| isoniazid_72h | tORF_34630 |
| isoniazid_72h | tORF_95238 |
| isoniazid_72h,pyranizoic_72h,rifampicin_72h | tORF_14178 |
| isoniazid_72h,pyranizoic_04h | tORF_26189 |
| isoniazid_72h,pyranizoic_72h | tORF_30261 |
| isoniazid_72h,rifampicin_72h | tORF_35700 |
| isoniazid_72h,rifampicin_72h | tORF_67521 |
| isoniazid_72h,pyranizoic_24h,pyranizoic_72h,rifampicin_04h | tORF_72141 |
| isoniazid_72h,rifampicin_24h,rifampicin_72h | tORF_79117 |
| isoniazid_72h,rifampicin_24h,rifampicin_72h | tORF_87534 |
| isoniazid_72h,rifampicin_24h,rifampicin_72h | tORF_103888 |
| isoniazid_72h,rifampicin_04h,rifampicin_72h | tORF_92637 |
| pyranizoic_24h | tORF_28759 |
| pyranizoic_72h | tORF_30521 |

| | |
|---|---|
| pyranizoic_72h,rifampicin_72h | tORF_45152 |
| rifampicin_24h,rifampicin_72h | tORF_11163 |
| rifampicin_24h,rifampicin_72h | tORF_17297 |
| rifampicin_24h,rifampicin_72h | tORF_33916 |
| rifampicin_24h,rifampicin_72h | tORF_53372 |
| rifampicin_24h | tORFs_11242_11233 |
| rifampicin_24h | tORF_49271 |
| rifampicin_24h | tORF_67823 |
| rifampicin_04h | tORF_94315 |
| rifampicin_72h | tORF_3808 |
| rifampicin_72h | tORF_12232 |
| rifampicin_72h | tORF_35369 |
| rifampicin_72h | tORF_36126 |
| rifampicin_72h | tORF_38722 |
| rifampicin_72h | tORF_69156 |
| rifampicin_72h | tORF_69185 |
| rifampicin_72h | tORF_71284 |
| rifampicin_72h | tORF_80148 |
| rifampicin_72h | tORF_84912 |
| rifampicin_72h | tORF_99968 |

Supplementary Table 6. Differential expression analysis of smORFs during bacterial starvation. Available as a separate Excel file (see Supplementary Materials).

Supplementary Table 7. Inferring smORFs' function from network co-expression analysis. Available as a separate Excel file (see Supplementary Materials).

Supplementary Table 8: Gene Ontology terms for annotated genes that are co-expressed with at least one SCT.

| Term | Name | Ontology Domain |
|---|---|---|
| nitrogen utilization | Rv2220 | biological process |
| nitrogen utilization | Rv3859c | biological process |
| zinc ion homeostasis | Rv0282 | biological process |
| zinc ion homeostasis | Rv2025c | biological process |
| divalent inorganic cation homeostasis | Rv0282 | biological process |
| divalent inorganic cation homeostasis | Rv2025c | biological process |
| RNA (guanine-N7)-methylation | Rv0208c | biological process |
| RNA (guanine-N7)-methylation | Rv3919c | biological process |
| daunorubicin transport | Rv2938 | biological process |
| daunorubicin transport | Rv2937 | biological process |
| daunorubicin transport | Rv2936 | biological process |
| glycoside transport | Rv2938 | biological process |
| glycoside transport | Rv2937 | biological process |
| glycoside transport | Rv2936 | biological process |
| doxorubicin transport | Rv2938 | biological process |
| doxorubicin transport | Rv2937 | biological process |
| doxorubicin transport | Rv2936 | biological process |
| organic hydroxy compound transport | Rv2938 | biological process |
| organic hydroxy compound transport | Rv2937 | biological process |
| organic hydroxy compound transport | Rv2936 | biological process |
| iron-sulfur cluster assembly | Rv1462 | biological process |
| iron-sulfur cluster assembly | Rv1461 | biological process |
| iron-sulfur cluster assembly | Rv1465 | biological process |
| iron-sulfur cluster assembly | Rv2204c | biological process |
| metallo-sulfur cluster assembly | Rv1462 | biological process |
| metallo-sulfur cluster assembly | Rv1461 | biological process |
| metallo-sulfur cluster assembly | Rv1465 | biological process |
| metallo-sulfur cluster assembly | Rv2204c | biological process |
| rRNA base methylation | Rv2372c | biological process |
| rRNA base methylation | Rv2165c | biological process |
| rRNA base methylation | Rv3919c | biological process |
| ketone biosynthetic process | Rv0534c | biological process |
| ketone biosynthetic process | Rv1372 | biological process |
| ketone biosynthetic process | Rv0542c | biological process |

| | | |
|---|---|---|
| ketone biosynthetic process | Rv0558 | biological process |
| cellular ketone metabolic process | Rv0534c | biological process |
| cellular ketone metabolic process | Rv1372 | biological process |
| cellular ketone metabolic process | Rv0542c | biological process |
| cellular ketone metabolic process | Rv1716 | biological process |
| cellular ketone metabolic process | Rv0558 | biological process |
| nitrate metabolic process | Rv1736c | biological process |
| nitrate metabolic process | Rv0818 | biological process |
| nitrate metabolic process | Rv1162 | biological process |
| nitrate metabolic process | Rv0267 | biological process |
| response to metal ion | Rv2031c | biological process |
| response to metal ion | Rv3224 | biological process |
| response to metal ion | Rv1994c | biological process |
| response to metal ion | Rv2642 | biological process |
| response to metal ion | Rv2641 | biological process |
| response to metal ion | Rv1992c | biological process |
| response to metal ion | Rv1909c | biological process |
| response to inorganic substance | Rv2031c | biological process |
| response to inorganic substance | Rv3224 | biological process |
| response to inorganic substance | Rv1162 | biological process |
| response to inorganic substance | Rv1994c | biological process |
| response to inorganic substance | Rv3130c | biological process |
| response to inorganic substance | Rv2642 | biological process |
| response to inorganic substance | Rv2641 | biological process |
| response to inorganic substance | Rv1992c | biological process |
| response to inorganic substance | Rv1760 | biological process |
| response to inorganic substance | Rv1221 | biological process |
| response to inorganic substance | Rv1909c | biological process |
| response to inorganic substance | Rv3234c | biological process |
| response to chemical | Rv3224 | biological process |
| response to chemical | Rv0821c | biological process |
| response to chemical | Rv1162 | biological process |
| response to chemical | Rv3286c | biological process |
| response to chemical | Rv0596c | biological process |
| response to chemical | Rv3161c | biological process |
| response to chemical | Rv3171c | biological process |
| response to chemical | Rv1760 | biological process |
| response to chemical | Rv1221 | biological process |
| response to chemical | Rv2238c | biological process |
| response to chemical | Rv1767 | biological process |
| response to chemical | Rv1909c | biological process |
| response to chemical | Rv2937 | biological process |

| | | |
|---|---|---|
| response to chemical | Rv2878c | biological process |
| response to chemical | Rv2936 | biological process |
| response to chemical | Rv3526 | biological process |
| response to chemical | Rv2258c | biological process |
| response to chemical | Rv1988 | biological process |
| response to chemical | Rv2031c | biological process |
| response to chemical | Rv2938 | biological process |
| response to chemical | Rv1218c | biological process |
| response to chemical | Rv0262c | biological process |
| response to chemical | Rv3104c | biological process |
| response to chemical | Rv3132c | biological process |
| response to chemical | Rv3130c | biological process |
| response to chemical | Rv1531 | biological process |
| response to chemical | Rv2642 | biological process |
| response to chemical | Rv2641 | biological process |
| response to chemical | Rv0342 | biological process |
| response to chemical | Rv3160c | biological process |
| response to chemical | Rv1536 | biological process |
| response to chemical | Rv0983 | biological process |
| response to chemical | Rv2005c | biological process |
| response to chemical | Rv0888 | biological process |
| response to chemical | Rv1955 | biological process |
| response to chemical | Rv1994c | biological process |
| response to chemical | Rv1991A | biological process |
| response to chemical | Rv2030c | biological process |
| response to chemical | Rv1992c | biological process |
| response to chemical | Rv3854c | biological process |
| response to chemical | Rv1217c | biological process |
| response to chemical | Rv3234c | biological process |
| response to stimulus | Rv0798c | biological process |
| response to stimulus | Rv1462 | biological process |
| response to stimulus | Rv3880c | biological process |
| response to stimulus | Rv2816c | biological process |
| response to stimulus | Rv3343c | biological process |
| response to stimulus | Rv2032 | biological process |
| response to stimulus | Rv3286c | biological process |
| response to stimulus | Rv2626c | biological process |
| response to stimulus | Rv0596c | biological process |
| response to stimulus | Rv0891c | biological process |
| response to stimulus | Rv3161c | biological process |
| response to stimulus | Rv1221 | biological process |
| response to stimulus | Rv2238c | biological process |

| | | |
|---|---|---|
| response to stimulus | Rv1821 | biological process |
| response to stimulus | Rv1909c | biological process |
| response to stimulus | Rv3645 | biological process |
| response to stimulus | Rv3526 | biological process |
| response to stimulus | Rv2258c | biological process |
| response to stimulus | Rv1988 | biological process |
| response to stimulus | Rv2031c | biological process |
| response to stimulus | Rv0818 | biological process |
| response to stimulus | Rv1218c | biological process |
| response to stimulus | Rv0042c | biological process |
| response to stimulus | Rv2051c | biological process |
| response to stimulus | Rv3134c | biological process |
| response to stimulus | Rv3130c | biological process |
| response to stimulus | Rv3571 | biological process |
| response to stimulus | Rv2642 | biological process |
| response to stimulus | Rv2641 | biological process |
| response to stimulus | Rv0342 | biological process |
| response to stimulus | Rv2887 | biological process |
| response to stimulus | Rv0983 | biological process |
| response to stimulus | Rv2007c | biological process |
| response to stimulus | Rv1955 | biological process |
| response to stimulus | Rv2373c | biological process |
| response to stimulus | Rv2030c | biological process |
| response to stimulus | Rv1992c | biological process |
| response to stimulus | Rv1027c | biological process |
| response to stimulus | Rv3854c | biological process |
| response to stimulus | Rv1217c | biological process |
| response to stimulus | Rv3765c | biological process |
| response to stimulus | Rv3224 | biological process |
| response to stimulus | Rv0821c | biological process |
| response to stimulus | Rv1162 | biological process |
| response to stimulus | Rv2212 | biological process |
| response to stimulus | Rv3171c | biological process |
| response to stimulus | Rv1760 | biological process |
| response to stimulus | Rv1767 | biological process |
| response to stimulus | Rv2937 | biological process |
| response to stimulus | Rv2503c | biological process |
| response to stimulus | Rv2878c | biological process |
| response to stimulus | Rv2936 | biological process |
| response to stimulus | Rv2938 | biological process |
| response to stimulus | Rv2110c | biological process |
| response to stimulus | Rv0918 | biological process |

| | | |
|---|---|---|
| response to stimulus | Rv0262c | biological process |
| response to stimulus | Rv3104c | biological process |
| response to stimulus | Rv0282 | biological process |
| response to stimulus | Rv3132c | biological process |
| response to stimulus | Rv3875 | biological process |
| response to stimulus | Rv1531 | biological process |
| response to stimulus | Rv1696 | biological process |
| response to stimulus | Rv3160c | biological process |
| response to stimulus | Rv1536 | biological process |
| response to stimulus | Rv0264c | biological process |
| response to stimulus | Rv2623 | biological process |
| response to stimulus | Rv2941 | biological process |
| response to stimulus | Rv2629 | biological process |
| response to stimulus | Rv1738 | biological process |
| response to stimulus | Rv2005c | biological process |
| response to stimulus | Rv0888 | biological process |
| response to stimulus | Rv1736c | biological process |
| response to stimulus | Rv1994c | biological process |
| response to stimulus | Rv1991A | biological process |
| response to stimulus | Rv3674c | biological process |
| response to stimulus | Rv3234c | biological process |
| response to hypoxia | Rv3134c | biological process |
| response to hypoxia | Rv2032 | biological process |
| response to hypoxia | Rv2626c | biological process |
| response to hypoxia | Rv3130c | biological process |
| response to hypoxia | Rv3571 | biological process |
| response to hypoxia | Rv1760 | biological process |
| response to hypoxia | Rv2623 | biological process |
| response to hypoxia | Rv2007c | biological process |
| response to hypoxia | Rv2629 | biological process |
| response to hypoxia | Rv2005c | biological process |
| response to hypoxia | Rv2503c | biological process |
| response to hypoxia | Rv2258c | biological process |
| response to hypoxia | Rv1955 | biological process |
| response to hypoxia | Rv2031c | biological process |
| response to hypoxia | Rv1736c | biological process |
| response to hypoxia | Rv3234c | biological process |
| response to decreased oxygen levels | Rv3134c | biological process |
| response to decreased oxygen levels | Rv2032 | biological process |
| response to decreased oxygen levels | Rv1162 | biological process |
| response to decreased oxygen levels | Rv3286c | biological process |
| response to decreased oxygen levels | Rv2626c | biological process |

| | | |
|---|---|---|
| response to decreased oxygen levels | Rv3130c | biological process |
| response to decreased oxygen levels | Rv3571 | biological process |
| response to decreased oxygen levels | Rv1760 | biological process |
| response to decreased oxygen levels | Rv2623 | biological process |
| response to decreased oxygen levels | Rv2007c | biological process |
| response to decreased oxygen levels | Rv2629 | biological process |
| response to decreased oxygen levels | Rv2005c | biological process |
| response to decreased oxygen levels | Rv2503c | biological process |
| response to decreased oxygen levels | Rv2258c | biological process |
| response to decreased oxygen levels | Rv1955 | biological process |
| response to decreased oxygen levels | Rv2031c | biological process |
| response to decreased oxygen levels | Rv1736c | biological process |
| response to decreased oxygen levels | Rv3234c | biological process |
| response to oxygen levels | Rv3134c | biological process |
| response to oxygen levels | Rv2032 | biological process |
| response to oxygen levels | Rv1162 | biological process |
| response to oxygen levels | Rv3286c | biological process |
| response to oxygen levels | Rv2626c | biological process |
| response to oxygen levels | Rv3130c | biological process |
| response to oxygen levels | Rv3571 | biological process |
| response to oxygen levels | Rv1760 | biological process |
| response to oxygen levels | Rv2623 | biological process |
| response to oxygen levels | Rv2007c | biological process |
| response to oxygen levels | Rv2629 | biological process |
| response to oxygen levels | Rv2005c | biological process |
| response to oxygen levels | Rv2503c | biological process |
| response to oxygen levels | Rv2258c | biological process |
| response to oxygen levels | Rv1955 | biological process |
| response to oxygen levels | Rv2031c | biological process |
| response to oxygen levels | Rv1736c | biological process |
| response to oxygen levels | Rv3234c | biological process |
| response to abiotic stimulus | Rv3134c | biological process |
| response to abiotic stimulus | Rv2032 | biological process |
| response to abiotic stimulus | Rv1162 | biological process |
| response to abiotic stimulus | Rv3286c | biological process |
| response to abiotic stimulus | Rv2626c | biological process |
| response to abiotic stimulus | Rv3130c | biological process |
| response to abiotic stimulus | Rv3571 | biological process |
| response to abiotic stimulus | Rv1696 | biological process |
| response to abiotic stimulus | Rv1760 | biological process |
| response to abiotic stimulus | Rv1221 | biological process |
| response to abiotic stimulus | Rv2623 | biological process |

| | | |
|---|---|---|
| response to abiotic stimulus | Rv2007c | biological process |
| response to abiotic stimulus | Rv2629 | biological process |
| response to abiotic stimulus | Rv2005c | biological process |
| response to abiotic stimulus | Rv2503c | biological process |
| response to abiotic stimulus | Rv2258c | biological process |
| response to abiotic stimulus | Rv1955 | biological process |
| response to abiotic stimulus | Rv2031c | biological process |
| response to abiotic stimulus | Rv2373c | biological process |
| response to abiotic stimulus | Rv1736c | biological process |
| response to abiotic stimulus | Rv3104c | biological process |
| response to abiotic stimulus | Rv3234c | biological process |
| DIM/DIP cell wall layer assembly | Rv2946c | biological process |
| DIM/DIP cell wall layer assembly | Rv2939 | biological process |
| DIM/DIP cell wall layer assembly | Rv2932 | biological process |
| DIM/DIP cell wall layer assembly | Rv2931 | biological process |
| DIM/DIP cell wall layer assembly | Rv2941 | biological process |
| DIM/DIP cell wall layer assembly | Rv2937 | biological process |
| DIM/DIP cell wall layer assembly | Rv2935 | biological process |
| DIM/DIP cell wall layer assembly | Rv2048c | biological process |
| cellular response to DNA damage stimulus | Rv1696 | biological process |
| cellular response to DNA damage stimulus | Rv3674c | biological process |
| plasma membrane fumarate reductase complex | Rv1554 | cellular component |
| plasma membrane fumarate reductase complex | Rv1553 | cellular component |
| plasma membrane fumarate reductase complex | Rv1555 | cellular component |
| fumarate reductase complex | Rv1554 | cellular component |
| fumarate reductase complex | Rv1553 | cellular component |
| fumarate reductase complex | Rv1555 | cellular component |
| respiratory chain complex II | Rv1554 | cellular component |
| respiratory chain complex II | Rv1553 | cellular component |
| respiratory chain complex II | Rv1555 | cellular component |
| plasma membrane respiratory chain complex II | Rv1554 | cellular component |
| plasma membrane respiratory chain complex II | Rv1553 | cellular component |
| plasma membrane respiratory chain complex II | Rv1555 | cellular component |
| intracellular non-membrane-bounded organelle | Rv0716 | cellular component |
| intracellular non-membrane-bounded organelle | Rv1630 | cellular component |
| intracellular organelle | Rv0716 | cellular component |

| | | |
|---|---|---|
| intracellular organelle | Rv0884c | cellular component |
| intracellular organelle | Rv1630 | cellular component |
| organelle | Rv0716 | cellular component |
| organelle | Rv0884c | cellular component |
| organelle | Rv1630 | cellular component |
| non-membrane-bounded organelle | Rv0716 | cellular component |
| non-membrane-bounded organelle | Rv1630 | cellular component |
| 3-oxoacid CoA-transferase activity | Rv2504c | molecular function |
| 3-oxoacid CoA-transferase activity | Rv2503c | molecular function |
| 3-ketosteroid 9-alpha-monooxygenase activity | Rv3571 | molecular function |
| 3-ketosteroid 9-alpha-monooxygenase activity | Rv3526 | molecular function |
| oxidoreductase activity, acting on paired donors, with incorporation or reduction of molecular oxygen, NAD(P)H as one donor, and incorporation of one atom of oxygen | Rv3518c | molecular function |
| oxidoreductase activity, acting on paired donors, with incorporation or reduction of molecular oxygen, NAD(P)H as one donor, and incorporation of one atom of oxygen | Rv3571 | molecular function |
| oxidoreductase activity, acting on paired donors, with incorporation or reduction of molecular oxygen, NAD(P)H as one donor, and incorporation of one atom of oxygen | Rv2266 | molecular function |
| oxidoreductase activity, acting on paired donors, with incorporation or reduction of molecular oxygen, NAD(P)H as one donor, and incorporation of one atom of oxygen | Rv3854c | molecular function |
| oxidoreductase activity, acting on paired donors, with incorporation or reduction of molecular oxygen, NAD(P)H as one donor, and incorporation of one atom of oxygen | Rv3049c | molecular function |
| oxidoreductase activity, acting on paired donors, with incorporation or reduction of molecular oxygen, NAD(P)H as one donor, and incorporation of one atom of oxygen | Rv3526 | molecular function |
| ketosteroid monooxygenase activity | Rv3571 | molecular function |
| ketosteroid monooxygenase activity | Rv3526 | molecular function |
| acetate CoA-transferase activity | Rv2504c | molecular function |
| acetate CoA-transferase activity | Rv2503c | molecular function |
| 3 iron, 4 sulfur cluster binding | Rv3859c | molecular function |
| 3 iron, 4 sulfur cluster binding | Rv1553 | molecular function |
| 3 iron, 4 sulfur cluster binding | Rv2007c | molecular function |
| iron-sulfur cluster binding | Rv2868c | molecular function |
| iron-sulfur cluster binding | Rv3138 | molecular function |
| iron-sulfur cluster binding | Rv3571 | molecular function |

| | | |
|---|---|---|
| iron-sulfur cluster binding | Rv1553 | molecular function |
| iron-sulfur cluster binding | Rv1465 | molecular function |
| iron-sulfur cluster binding | Rv3161c | molecular function |
| iron-sulfur cluster binding | Rv3260c | molecular function |
| iron-sulfur cluster binding | Rv2218 | molecular function |
| iron-sulfur cluster binding | Rv0886 | molecular function |
| iron-sulfur cluster binding | Rv2204c | molecular function |
| iron-sulfur cluster binding | Rv2007c | molecular function |
| iron-sulfur cluster binding | Rv3526 | molecular function |
| iron-sulfur cluster binding | Rv2900c | molecular function |
| iron-sulfur cluster binding | Rv1736c | molecular function |
| iron-sulfur cluster binding | Rv3859c | molecular function |
| iron-sulfur cluster binding | Rv3503c | molecular function |
| iron-sulfur cluster binding | Rv3674c | molecular function |
| iron-sulfur cluster binding | Rv3862c | molecular function |
| metal cluster binding | Rv2868c | molecular function |
| metal cluster binding | Rv3138 | molecular function |
| metal cluster binding | Rv3571 | molecular function |
| metal cluster binding | Rv1553 | molecular function |
| metal cluster binding | Rv1465 | molecular function |
| metal cluster binding | Rv3161c | molecular function |
| metal cluster binding | Rv3260c | molecular function |
| metal cluster binding | Rv2218 | molecular function |
| metal cluster binding | Rv0886 | molecular function |
| metal cluster binding | Rv2204c | molecular function |
| metal cluster binding | Rv2007c | molecular function |
| metal cluster binding | Rv3526 | molecular function |
| metal cluster binding | Rv2900c | molecular function |
| metal cluster binding | Rv1736c | molecular function |
| metal cluster binding | Rv3859c | molecular function |
| metal cluster binding | Rv3503c | molecular function |
| metal cluster binding | Rv3674c | molecular function |
| metal cluster binding | Rv3862c | molecular function |
| N-methyltransferase activity | Rv2372c | molecular function |
| N-methyltransferase activity | Rv2165c | molecular function |
| N-methyltransferase activity | Rv3919c | molecular function |
| N-methyltransferase activity | Rv1988 | molecular function |
| oxidoreductase activity, acting on the CH-NH group of donors, NAD or NADP as acceptor | Rv1187 | molecular function |
| oxidoreductase activity, acting on the CH-NH group of donors, NAD or NADP as acceptor | Rv0245 | molecular function |
| oxidoreductase activity, acting on the CH-NH group of donors, NAD or NADP as acceptor | Rv0500 | molecular function |

| | | |
|---|---|---|
| oxidoreductase activity, acting on the CH-NH group of donors, NAD or NADP as acceptor | Rv3007c | molecular function |
| oxidoreductase activity, acting on the CH-NH group of donors | Rv1187 | molecular function |
| oxidoreductase activity, acting on the CH-NH group of donors | Rv0245 | molecular function |
| oxidoreductase activity, acting on the CH-NH group of donors | Rv0500 | molecular function |
| oxidoreductase activity, acting on the CH-NH group of donors | Rv3007c | molecular function |
| rRNA methyltransferase activity | Rv2372c | molecular function |
| rRNA methyltransferase activity | Rv2165c | molecular function |
| rRNA methyltransferase activity | Rv3919c | molecular function |
| rRNA methyltransferase activity | Rv1988 | molecular function |
| catalytic activity, acting on a rRNA | Rv2372c | molecular function |
| catalytic activity, acting on a rRNA | Rv2165c | molecular function |
| catalytic activity, acting on a rRNA | Rv3919c | molecular function |
| catalytic activity, acting on a rRNA | Rv1988 | molecular function |
| metalloendopeptidase activity | Rv2869c | molecular function |
| metalloendopeptidase activity | Rv0563 | molecular function |
| metalloendopeptidase activity | Rv2782c | molecular function |
| metalloendopeptidase activity | Rv2625c | molecular function |
| metalloendopeptidase activity | Rv1977 | molecular function |
| 2 iron, 2 sulfur cluster binding | Rv3571 | molecular function |
| 2 iron, 2 sulfur cluster binding | Rv1553 | molecular function |
| 2 iron, 2 sulfur cluster binding | Rv1465 | molecular function |
| 2 iron, 2 sulfur cluster binding | Rv3161c | molecular function |
| 2 iron, 2 sulfur cluster binding | Rv2204c | molecular function |
| 2 iron, 2 sulfur cluster binding | Rv3526 | molecular function |
| transaminase activity | Rv0884c | molecular function |
| transaminase activity | Rv1178 | molecular function |
| transaminase activity | Rv3772 | molecular function |
| transaminase activity | Rv3290c | molecular function |
| transaminase activity | Rv1005c | molecular function |
| transaminase activity | Rv3436c | molecular function |
| transaminase activity | Rv3329 | molecular function |
| transferase activity, transferring nitrogenous groups | Rv0884c | molecular function |
| transferase activity, transferring nitrogenous groups | Rv1178 | molecular function |
| transferase activity, transferring nitrogenous groups | Rv3772 | molecular function |
| transferase activity, transferring nitrogenous groups | Rv3290c | molecular function |

| | | |
|---|---|---|
| transferase activity, transferring nitrogenous groups | Rv1005c | molecular function |
| transferase activity, transferring nitrogenous groups | Rv3436c | molecular function |
| transferase activity, transferring nitrogenous groups | Rv3329 | molecular function |
| 4 iron, 4 sulfur cluster binding | Rv1736c | molecular function |
| 4 iron, 4 sulfur cluster binding | Rv2868c | molecular function |
| 4 iron, 4 sulfur cluster binding | Rv3138 | molecular function |
| 4 iron, 4 sulfur cluster binding | Rv1553 | molecular function |
| 4 iron, 4 sulfur cluster binding | Rv3260c | molecular function |
| 4 iron, 4 sulfur cluster binding | Rv2218 | molecular function |
| 4 iron, 4 sulfur cluster binding | Rv0886 | molecular function |
| 4 iron, 4 sulfur cluster binding | Rv2204c | molecular function |
| 4 iron, 4 sulfur cluster binding | Rv2007c | molecular function |
| 4 iron, 4 sulfur cluster binding | Rv3674c | molecular function |
| 4 iron, 4 sulfur cluster binding | Rv3862c | molecular function |
| 4 iron, 4 sulfur cluster binding | Rv2900c | molecular function |
| catalytic activity, acting on DNA | Rv0797 | molecular function |
| catalytic activity, acting on DNA | Rv3674c | molecular function |

Supplementary Table 9: smORFs homologs found during blastp and tblastn searches

| smorf | blastp actinobacteria | blastp mycobacteria | tblastn actinobacteria | tblastn mycobacteria |
|---|---|---|---|---|
| tORF_47070 | 0 | 0 | 0 | 22 |
| tORF_35175 | 0 | 0 | 51 | 100 |
| tORF_26369 | 0 | 0 | 0 | 100 |
| tORF_60096 | 0 | 0 | 0 | 104 |
| tORF_79117 | 0 | 3 | 0 | 59 |
| tORF_61297 | 0 | 0 | 0 | 29 |
| tORF_19112 | 0 | 0 | 100 | 100 |
| tORF_103888 | 0 | 0 | 0 | 59 |
| tORF_38722 | 0 | 0 | 0 | 6 |
| tORF_28759 | 0 | 0 | 0 | 108 |
| tORF_69156 | 0 | 0 | 0 | 6 |
| tORF_35700 | 0 | 0 | 0 | 103 |
| tORF_11242 | 0 | 0 | 69 | 101 |
| tORF_33916 | 0 | 0 | 1 | 100 |
| tORF_2047 | 0 | 0 | 0 | 4 |
| tORF_72141 | 0 | 0 | 0 | 44 |
| tORF_42121 | 0 | 0 | 0 | 0 |
| tORF_94745 | 0 | 0 | 0 | 1 |
| gORF_25784 | 0 | 0 | 2 | 28 |
| gORF_36517 | 100 | 100 | 100 | 100 |
| gORF_74745 | 0 | 0 | 0 | 51 |
| gORF_113929 | 0 | 13 | 0 | 258 |
| gORF_69007 | 0 | 0 | 0 | 152 |
| gORF_69072 | 0 | 0 | 0 | 22 |
| gORF_20316 | 0 | 1 | 14 | 100 |
| gORF_101064 | 0 | 0 | 0 | 1 |
| gORF_35414 | 0 | 5 | 105 | 136 |
| gORF_68556 | 0 | 5 | 96 | 100 |
| gORF_103397 | 0 | 22 | 0 | 18 |
| gORF_105604 | 0 | 0 | 0 | 6 |
| gORF_78109 | 0 | 0 | 1 | 100 |
| gORF_74686 | 0 | 0 | 0 | 0 |
| gORF_121862 | 0 | 2 | 103 | 100 |

| | | | | |
|---|---|---|---|---|
| gORF_92020 | 0 | 1 | 0 | 100 |
| gORF_37048 | 0 | 1 | 33 | 95 |
| gORF_66553 | 0 | 1 | 106 | 213 |
| gORF_75976 | 0 | 0 | 0 | 0 |
| gORF_35757 | 0 | 0 | 0 | 4 |
| gORF_90691 | 0 | 0 | 0 | 3 |
| gORF_6673 | 0 | 0 | 18 | 100 |
| gORF_107229 | 0 | 1 | 0 | 15 |
| gORF_32165 | 0 | 0 | 2 | 53 |
| gORF_2002 | 0 | 0 | 0 | 100 |
| gORF_7579 | 0 | 1 | 6 | 103 |
| tORF_92272 | 0 | 0 | 0 | 30 |
| tORF_100857 | 0 | 0 | 0 | 100 |
| tORF_88201 | 0 | 0 | 0 | 13 |
| tORF_56419 | 0 | 0 | 0 | 6 |
| tORF_99968 | 5 | 5 | 101 | 101 |
| tORF_87698 | 0 | 2 | 0 | 100 |
| tORF_89298 | 0 | 0 | 1 | 101 |
| tORF_75796 | 0 | 0 | 0 | 5 |
| gORF_68833 | 0 | 1 | 0 | 101 |
| gORF_57646 | 0 | 0 | 0 | 6 |
| gORF_50161 | 0 | 0 | 0 | 11 |
| gORF_106483 | 0 | 0 | 2 | 100 |
| gORF_48161 | 0 | 0 | 0 | 21 |
| gORF_42930 | 0 | 0 | 0 | 11 |
| gORF_11817 | 0 | 0 | 0 | 11 |
| gORF_47860 | 1 | 1 | 117 | 132 |
| gORF_29356 | 0 | 0 | 0 | 8 |
| gORF_95760 | 0 | 0 | 0 | 6 |
| gORF_75871 | 0 | 0 | 10 | 23 |
| tORF_288 | 0 | 1 | 20 | 100 |
| tORF_55810 | 0 | 0 | 0 | 0 |
| tORF_67857 | 0 | 0 | 0 | 5 |
| tORF_10760 | 0 | 0 | 0 | 421 |
| tORF_67521 | 0 | 0 | 0 | 100 |
| tORF_49343 | 0 | 0 | 0 | 10 |
| tORF_92637 | 0 | 0 | 0 | 4 |
| tORF_24534 | 0 | 0 | 0 | 5 |
| tORF_30521 | 0 | 4 | 104 | 125 |
| tORF_102211 | 0 | 0 | 0 | 0 |
| tORF_82961 | 0 | 1 | 0 | 100 |
| tORF_76742 | 0 | 0 | 0 | 63 |

| | | | | |
|---|---|---|---|---|
| tORF_103255 | 0 | 0 | 0 | 37 |
| tORF_72971 | 0 | 1 | 0 | 100 |
| tORF_36024 | 0 | 0 | 0 | 6 |
| tORF_14178 | 0 | 0 | 0 | 7 |
| tORF_22610 | 0 | 0 | 7 | 99 |
| tORF_9636 | 0 | 0 | 0 | 5 |
| tORF_53372 | 0 | 0 | 1 | 44 |
| tORF_34630 | 0 | 0 | 0 | 15 |
| tORF_29093 | 0 | 0 | 108 | 112 |
| tORF_12940 | 0 | 0 | 14 | 101 |
| tORF_4175 | 0 | 0 | 0 | 34 |
| tORF_89943 | 0 | 0 | 0 | 4 |
| tORF_11163 | 0 | 0 | 0 | 26 |
| tORF_88334 | 0 | 0 | 0 | 130 |
| tORF_67974 | 0 | 0 | 0 | 4 |
| tORF_20207 | 0 | 1 | 0 | 7 |
| tORF_45152 | 0 | 0 | 1 | 87 |
| tORF_33963 | 0 | 0 | 0 | 14 |
| tORF_90234 | 0 | 1 | 24 | 108 |
| tORF_8883 | 0 | 0 | 0 | 8 |
| tORF_22787 | 0 | 0 | 0 | 119 |
| tORF_66504 | 0 | 4 | 0 | 71 |
| tORF_17297 | 0 | 0 | 0 | 8 |
| tORF_94315 | 0 | 0 | 0 | 93 |
| tORF_87534 | 0 | 0 | 19 | 14 |
| tORF_90137 | 0 | 0 | 0 | 7 |
| tORF_89037 | 0 | 1 | 62 | 100 |
| tORF_21472 | 0 | 1 | 3 | 100 |
| tORF_20553 | 0 | 0 | 0 | 0 |
| tORF_65674 | 0 | 0 | 0 | 28 |
| tORF_438 | 0 | 1 | 0 | 100 |
| tORF_437 | 0 | 1 | 0 | 100 |
| tORF_433 | 0 | 1 | 1 | 100 |
| tORF_81913 | 0 | 0 | 46 | 100 |
| tORF_63427 | 3 | 0 | 100 | 100 |
| tORF_67941 | 0 | 0 | 0 | 6 |
| tORF_9734 | 0 | 0 | 0 | 82 |
| tORF_98881 | 0 | 0 | 4 | 100 |
| tORF_4671 | 0 | 0 | 0 | 0 |
| tORF_92330 | 0 | 0 | 0 | 83 |
| tORF_38551 | 0 | 0 | 0 | 59 |
| tORF_39817 | 0 | 0 | 0 | 10 |

| | | | | |
|---|---|---|---|---|
| tORF_39816 | 0 | 0 | 0 | 10 |
| tORF_16103 | 0 | 0 | 0 | 0 |
| tORF_26465 | 0 | 1 | 0 | 6 |
| tORF_8639 | 0 | 4 | 0 | 4 |
| tORF_104672 | 0 | 0 | 0 | 6 |
| tORF_7075 | 0 | 0 | 0 | 25 |
| tORF_39731 | 0 | 0 | 13 | 100 |
| tORF_29123 | 0 | 0 | 15 | 100 |
| tORF_69957 | 0 | 1 | 10 | 100 |
| tORF_74872 | 0 | 0 | 0 | 100 |
| tORF_69185 | 0 | 0 | 0 | 6 |
| tORF_30438 | 0 | 0 | 0 | 11 |
| tORF_11233 | 0 | 0 | 7 | 101 |
| tORF_3808 | 0 | 0 | 1 | 100 |
| tORF_47766 | 0 | 0 | 0 | 100 |
| tORF_60406 | 0 | 1 | 0 | 82 |
| tORF_26189 | 0 | 0 | 0 | 0 |
| tORF_104870 | 0 | 0 | 0 | 4 |
| tORF_71284 | 0 | 0 | 0 | 8 |
| tORF_84082 | 0 | 0 | 0 | 6 |
| tORF_29232 | 0 | 0 | 0 | 77 |
| tORF_84912 | 0 | 0 | 0 | 7 |
| tORF_73702 | 0 | 0 | 100 | 114 |
| tORF_63374 | 0 | 0 | 0 | 5 |
| tORF_68868 | 0 | 1 | 101 | 100 |
| tORF_30261 | 0 | 0 | 0 | 100 |
| tORF_76226 | 0 | 0 | 0 | 56 |
| tORF_12232 | 0 | 0 | 6 | 13 |
| tORF_49271 | 0 | 0 | 0 | 5 |
| tORF_102203 | 0 | 0 | 0 | 100 |
| tORF_74194 | 0 | 0 | 0 | 85 |
| tORF_47960 | 0 | 0 | 0 | 10 |
| tORF_3622 | 0 | 67 | 0 | 46 |
| tORF_67823 | 0 | 0 | 0 | 8 |
| tORF_421 | 0 | 0 | 0 | 100 |
| tORF_36126 | 0 | 0 | 0 | 6 |
| tORF_37787 | 0 | 0 | 0 | 12 |
| tORF_3869 | 0 | 0 | 0 | 2 |
| tORF_80148 | 0 | 0 | 0 | 43 |
| tORF_53851 | 0 | 0 | 0 | 7 |
| tORF_88156 | 0 | 0 | 0 | 100 |
| tORF_73986 | 0 | 0 | 0 | 10 |

| | | | | |
|---|---|---|---|---|
| tORF_70947 | 0 | 0 | 0 | 0 |
| tORF_3074 | 0 | 8 | 3 | 732 |
| tORF_35369 | 0 | 0 | 0 | 6 |
| tORF_22109 | 0 | 0 | 0 | 6 |
| tORF_95238 | 0 | 0 | 0 | 100 |
| tORF_67492 | 0 | 0 | 2 | 100 |
| gORF_75773 | 0 | 0 | 0 | 51 |
| gORF_65795 | 0 | 0 | 0 | 26 |
| gORF_67446 | 0 | 4 | 27 | 104 |
| gORF_33946 | 0 | 0 | 0 | 46 |
| gORF_250 | 0 | 1 | 0 | 94 |
| gORF_59194 | 0 | 0 | 0 | 0 |
| gORF_48905 | 0 | 2 | 0 | 100 |
| gORF_1756 | 0 | 0 | 2 | 102 |
| gORF_41235 | 0 | 1 | 0 | 100 |
| gORF_51888 | 0 | 0 | 42 | 55 |
| gORF_5914 | 0 | 0 | 0 | 6 |
| gORF_85060 | 0 | 0 | 0 | 65 |
| gORF_65171 | 0 | 1 | 3 | 171 |
| gORF_120753 | 0 | 0 | 0 | 104 |
| gORF_87937 | 0 | 0 | 0 | 63 |
| gORF_90956 | 0 | 0 | 0 | 13 |
| gORF_68832 | 0 | 1 | 0 | 101 |
| gORF_68829 | 0 | 1 | 0 | 101 |
| gORF_106326 | 0 | 0 | 0 | 13 |
| gORF_62933 | 0 | 0 | 0 | 7 |
| gORF_117472 | 0 | 0 | 104 | 107 |
| gORF_76198 | 0 | 1 | 76 | 100 |
| gORF_102062 | 0 | 0 | 0 | 29 |
| gORF_52031 | 0 | 0 | 0 | 6 |
| gORF_422 | 0 | 2 | 2 | 100 |
| gORF_118806 | 0 | 0 | 0 | 6 |
| gORF_99913 | 0 | 0 | 0 | 6 |
| gORF_15589 | 1 | 3 | 27 | 100 |
| gORF_61558 | 0 | 0 | 0 | 37 |
| gORF_88422 | 0 | 0 | 0 | 100 |
| gORF_13284 | 0 | 0 | 0 | 3 |
| gORF_61964 | 0 | 0 | 68 | 69 |
| gORF_99760 | 0 | 3 | 104 | 152 |
| gORF_52583 | 0 | 0 | 0 | 31 |
| gORF_30484 | 0 | 2 | 3 | 583 |
| gORF_85045 | 0 | 0 | 141 | 114 |

| | | | | |
|---|---|---|---|---|
| gORF_31439 | 0 | 0 | 0 | 5 |
| gORF_33915 | 0 | 0 | 0 | 25 |
| gORF_106880 | 0 | 4 | 0 | 103 |
| gORF_14611 | 0 | 0 | 0 | 100 |
| gORF_43305 | 0 | 0 | 0 | 6 |
| gORF_84877 | 0 | 0 | 0 | 28 |
| gORF_42594 | 0 | 0 | 0 | 100 |
| gORF_443 | 0 | 0 | 0 | 6 |
| gORF_25839 | 0 | 0 | 0 | 6 |
| gORF_329 | 0 | 0 | 0 | 54 |
| gORF_3585 | 0 | 0 | 0 | 101 |
| gORF_118619 | 1 | 100 | 0 | 100 |
| gORF_80264 | 0 | 0 | 0 | 4 |
| gORF_102158 | 0 | 0 | 0 | 47 |
| gORF_1601 | 0 | 0 | 0 | 100 |
| gORF_92693 | 0 | 0 | 0 | 14 |
| gORF_107491 | 0 | 0 | 0 | 70 |
| gORF_9109 | 0 | 0 | 0 | 100 |
| gORF_82083 | 0 | 0 | 1 | 11 |
| gORF_19557 | 0 | 0 | 0 | 6 |
| gORF_61194 | 2 | 4 | 103 | 101 |
| gORF_100972 | 0 | 0 | 0 | 88 |
| gORF_61016 | 0 | 0 | 0 | 0 |
| gORF_65479 | 0 | 0 | 0 | 101 |
| gORF_119727 | 0 | 0 | 0 | 101 |
| gORF_14677 | 0 | 0 | 0 | 7 |
| gORF_7760 | 0 | 3 | 0 | 100 |
| gORF_53666 | 0 | 0 | 26 | 100 |
| gORF_101587 | 0 | 0 | 0 | 35 |
| gORF_33042 | 0 | 4 | 0 | 209 |
| gORF_29058 | 0 | 0 | 0 | 126 |
| gORF_48053 | 0 | 0 | 47 | 100 |
| gORF_28422 | 0 | 0 | 0 | 72 |
| gORF_104161 | 0 | 0 | 0 | 34 |
| gORF_87844 | 0 | 0 | 1 | 15 |
| gORF_41083 | 0 | 0 | 0 | 0 |
| gORF_46630 | 0 | 0 | 0 | 22 |
| gORF_119201 | 0 | 0 | 0 | 32 |
| gORF_104937 | 0 | 0 | 0 | 14 |
| gORF_38000 | 0 | 0 | 0 | 5 |
| gORF_376 | 0 | 0 | 0 | 100 |
| gORF_64858 | 0 | 1 | 0 | 100 |

| | | | | |
|---|---|---|---|---|
| gORF_2129 | 0 | 0 | 0 | 45 |
| gORF_80850 | 0 | 0 | 0 | 0 |
| gORF_806 | 0 | 0 | 0 | 6 |
| gORF_16021 | 0 | 1 | 0 | 29 |
| gORF_28867 | 0 | 1 | 1 | 100 |
| gORF_43180 | 0 | 0 | 7 | 43 |
| gORF_114282 | 1 | 1 | 100 | 100 |
| gORF_94851 | 0 | 1 | 0 | 6 |
| gORF_44038 | 0 | 3 | 129 | 100 |
| gORF_63052 | 0 | 0 | 0 | 7 |
| gORF_51269 | 0 | 0 | 153 | 102 |
| gORF_80350 | 0 | 0 | 0 | 6 |
| gORF_51571 | 0 | 72 | 0 | 1300 |
| gORF_57419 | 0 | 0 | 0 | 76 |
| gORF_68366 | 0 | 0 | 0 | 101 |
| gORF_34041 | 0 | 0 | 0 | 217 |
| gORF_111934 | 0 | 0 | 0 | 30 |
| gORF_84182 | 0 | 0 | 0 | 100 |
| gORF_62770 | 0 | 0 | 0 | 94 |
| gORF_35714 | 0 | 0 | 0 | 0 |
| gORF_28297 | 0 | 0 | 0 | 41 |
| gORF_91428 | 0 | 0 | 1 | 100 |
| gORF_84771 | 0 | 0 | 0 | 53 |
| gORF_56923 | 0 | 0 | 0 | 22 |
| gORF_5065 | 0 | 0 | 0 | 126 |
| gORF_7258 | 2 | 0 | 24 | 106 |
| gORF_40039 | 0 | 1 | 1 | 100 |
| gORF_74533 | 0 | 0 | 0 | 6 |
| gORF_46723 | 0 | 0 | 101 | 100 |
| gORF_44871 | 0 | 0 | 0 | 6 |
| gORF_106973 | 0 | 49 | 0 | 2049 |
| gORF_75042 | 0 | 0 | 0 | 6 |
| gORF_55538 | 0 | 0 | 0 | 4 |
| gORF_109100 | 0 | 1 | 0 | 127 |
| gORF_67285 | 1 | 3 | 927 | 247 |
| gORF_7303 | 0 | 0 | 0 | 56 |
| gORF_52560 | 0 | 0 | 0 | 9 |
| gORF_32818 | 0 | 0 | 0 | 100 |
| gORF_70944 | 0 | 0 | 9 | 126 |
| gORF_107153 | 0 | 1 | 91 | 101 |
| gORF_83262 | 0 | 0 | 18 | 102 |
| gORF_72705 | 0 | 0 | 0 | 100 |

| | | | | |
|---|---|---|---|---|
| gORF_19087 | 0 | 1 | 15 | 101 |
| gORF_50213 | 0 | 0 | 0 | 6 |
| gORF_27483 | 0 | 1 | 0 | 7 |
| gORF_114887 | 0 | 0 | 0 | 59 |
| gORF_105155 | 0 | 1 | 3 | 115 |
| gORF_2540 | 0 | 1 | 0 | 47 |
| gORF_19021 | 0 | 0 | 0 | 6 |
| gORF_113073 | 0 | 0 | 0 | 246 |
| gORF_95424 | 0 | 0 | 0 | 14 |
| gORF_102652 | 0 | 0 | 6 | 102 |
| gORF_105261 | 0 | 0 | 54 | 100 |
| gORF_86692 | 0 | 0 | 8 | 100 |
| gORF_38811 | 0 | 0 | 0 | 10 |
| gORF_105910 | 0 | 0 | 0 | 4 |
| gORF_68412 | 0 | 0 | 0 | 92 |
| gORF_42096 | 0 | 0 | 0 | 6 |
| gORF_117748 | 0 | 0 | 1 | 27 |
| gORF_101684 | 0 | 0 | 2 | 100 |
| gORF_85603 | 0 | 0 | 0 | 34 |
| gORF_68418 | 0 | 0 | 0 | 0 |
| gORF_52566 | 0 | 0 | 0 | 0 |
| gORF_112725 | 0 | 0 | 0 | 0 |
| gORF_67918 | 0 | 0 | 0 | 0 |
| gORF_38583 | 0 | 0 | 0 | 0 |
| gORF_78990 | 0 | 3 | 0 | 0 |
| gORF_67977 | 0 | 0 | 0 | 0 |
| gORF_55260 | 0 | 0 | 0 | 0 |
| gORF_45994 | 0 | 0 | 0 | 0 |
| gORF_67145 | 0 | 0 | 0 | 0 |
| gORF_18752 | 0 | 1 | 0 | 0 |
| gORF_105783 | 0 | 1 | 0 | 0 |
| gORF_72140 | 0 | 2 | 0 | 0 |
| gORF_120720 | 1 | 12 | 0 | 0 |
| gORF_27035 | 0 | 0 | 0 | 0 |
| gORF_77518 | 0 | 0 | 0 | 0 |
| gORF_26055 | 0 | 0 | 0 | 0 |
| gORF_56439 | 0 | 0 | 0 | 0 |
| gORF_91014 | 0 | 6 | 0 | 0 |

Supplementary Table 10: Full genera for species in the tree.

| **genus** | **full_name** |
|---|---|
| Conexibacter | Conexibacter_woesei |
| Geodermatophilus | Geodermatophilus_obscurus |
| Frankia | Frankia_inefficax |
| Blastococcus | Blastococcus_saxobsidens |
| Actinoplanes | Actinoplanes_globisporus |
| Actinomadura | Actinomadura_madurae |
| Nocardioides | Nocardioides_sp._URHA0032 |
| Saccharothrix | Saccharothrix_syringae |
| Tetrasphaera | Tetrasphaera_australiensis |
| Streptomyces | Streptomyces_silaceus |
| Planomonospora | Planomonospora_sphaerica |
| Enemella | Enemella_evansiae |
| Amycolatopsis | Amycolatopsis_sp._YIM |
| Ornithinibacter | Ornithinibacter_aureus |
| Lentzea | Lentzea_alba |
| Phycicoccus | Phycicoccus_sp._HDW14 |
| Pseudonocardia | Pseudonocardia_xinjiangensis |
| Nonomuraea | Nonomuraea_endophytica |
| Actinophytocola | Actinophytocola_algeriensis |
| Tenggerimyces | Tenggerimyces_flavus |
| Cryptosporangium | Cryptosporangium_aurantiacum |
| Actinokineospora | Actinokineospora_auranticolor |
| Marmoricola | Marmoricola_mangrovicus |
| Spirilliplanes | Spirilliplanes_yamanashiensis |
| Mycobacterium | Mycobacterium_avium |
| Rhodococcus | Rhodococcus_erythropolis |
| Mycolicibacterium | Mycolicibacterium_smegmatis |
| Mycobacteroides | Mycobacteroides_abscessus |
| Corynebacterium | Corynebacterium_striatum |
| Mycolicibacter | Mycolicibacter_kumamotonensis |
| Bifidobacterium | Bifidobacterium_longum |
| Leifsonia | Leifsonia_xyli |

| | |
|---|---|
| Cutibacterium | Cutibacterium_acnes |
| Kocuria | Kocuria_rhizophila |
| Paenarthrobacter | Paenarthrobacter_aurescens |
| Acidothermus | Acidothermus_cellulolyticus |
| Salinispora | Salinispora_tropica |
| Kineococcus | Kineococcus_radiotolerans |
| Pseudarthrobacter | Pseudarthrobacter_chlorophenolicus |
| Beutenbergia | Beutenbergia_cavernae |
| Micrococcus | Micrococcus_luteus |
| Actinosynnema | Actinosynnema_mirum |
| Saccharomonospora | Saccharomonospora_viridis |
| Kytococcus | Kytococcus_sedentarius |
| Catenulispora | Catenulispora_acidiphila |
| Glutamicibacter | Glutamicibacter_nicotianae |
| Micromonospora | Micromonospora_sagamiensis |
| Saccharopolyspora | Saccharopolyspora_pogona |
| Microbacterium | Microbacterium_hominis |
| Rathayibacter | Rathayibacter_toxicus |
| Clavibacter | Clavibacter_zhangzhiyongii |
| Eggerthella | Eggerthella_lenta |
| Propionibacterium | Propionibacterium_freudenreichii |
| Jonesia | Jonesia_denitrificans |
| Thermobispora | Thermobispora_bispora |
| Cellulosimicrobium | Cellulosimicrobium_cellulans |
| Brevibacterium | Brevibacterium_casei |
| Thermobifida | Thermobifida_halotolerans |
| Thermomonospora | Thermomonospora_amylolytica |
| Cryobacterium | Cryobacterium_soli |
| Gryllotalpicola | Gryllotalpicola_protaetiae |
| Protaetiibacter | Protaetiibacter_intestinalis |
| Arthrobacter | Arthrobacter_sp._C1-1 |
| Agromyces | Agromyces_marinus |
| Cellulomonas | Cellulomonas_iranensis |
| Actinomyces | Actinomyces_radicidentis |
| Rothia | Rothia_kristinae |
| Janibacter | Janibacter_limosus |
| Sinomonas | Sinomonas_atrocyanea |
| Frondihabitans | Frondihabitans_sp._PAMC |
| Pengzhenrongella | Pengzhenrongella_sicca |
| Dermatophilus | Dermatophilus_congolensis |
| Kutzneria | Kutzneria_albida |
| Leucobacter | Leucobacter_luti |

| | |
|---|---|
| Curtobacterium | Curtobacterium_flaccumfaciens |
| Luteimicrobium | Luteimicrobium_xylanilyticum |
| Raineyella | Raineyella_fluvialis |
| Aeromicrobium | Aeromicrobium_yanjiei |
| Pseudactinotalea | Pseudactinotalea_sp._HY158 |
| Parolsenella | Parolsenella_catena |
| Dermacoccus | Dermacoccus_nishinomiyaensis |
| Agrococcus | Agrococcus_jejuensis |
| Auraticoccus | Auraticoccus_monumenti |
| Prauserella | Prauserella_marina |
| Allokutzneria | Allokutzneria_albata |
| Nocardiopsis | Nocardiopsis_dassonvillei |
| Kitasatospora | Kitasatospora_mediocidica |
| Actinopolyspora | Actinopolyspora_erythraea |
| Intrasporangium | Intrasporangium_calvum |
| Euzebya | Euzebya_pacifica |
| Ornithinimicrobium | Ornithinimicrobium_avium |
| Plantibacter | Plantibacter_flavus |
| Schaalia | Schaalia_odontolytica |
| Isoptericola | Isoptericola_variabilis |
| Rubrobacter | Rubrobacter_radiotolerans |
| Microlunatus | Microlunatus_elymi |
| Verrucosispora | Verrucosispora_sp._NA02020 |
| Glycomyces | Glycomyces_sp._TRM65418 |
| Modestobacter | Modestobacter_marinus |
| Arcanobacterium | Arcanobacterium_phocae |
| Sanguibacter | Sanguibacter_sp._HDW7 |
| Brevilactibacter | Brevilactibacter_coleopterorum |
| Natronoglycomyces | Natronoglycomyces_albus |
| Natronosporangium | Natronosporangium_hydrolyticum |
| Georgenia | Georgenia_sp._TF02-10 |
| Austwickia | Austwickia_chelonae |
| Pimelobacter | Pimelobacter_simplex |
| Trueperella | Trueperella_pyogenes |
| Xylanimonas | Xylanimonas_allomyrinae |
| Kibdelosporangium | Kibdelosporangium_sp._MJ126-NF4 |
| Mycetocola | Mycetocola_zhujimingii |
| Aquihabitans | Aquihabitans_sp._G128 |
| Ruania | Ruania_zhangjianzhongii |
| Dermabacter | Dermabacter_vaginalis |
| Neomicrococcus | Neomicrococcus_aestuarii |
| Actinoalloteichus | Actinoalloteichus_hymeniacidonis |

| | |
|---|---|
| Arachnia | Arachnia_propionica |
| Acidipropionibacterium | Acidipropionibacterium_acidipropionici |
| Streptacidiphilus | Streptacidiphilus_griseoplanus |
| Alloactinosynnema | Alloactinosynnema_sp._L-07 |
| Microcella | Microcella_alkaliphila |
| Miltoncostaea | Miltoncostaea_marina |
| Gephyromycinifex | Gephyromycinifex_aptenodytis |
| Serinicoccus | Serinicoccus_hydrothermalis |
| Flaviflexus | Flaviflexus_ciconiae |
| Cnuibacter | Cnuibacter_physcomitrellae |
| Plantactinospora | Plantactinospora_sp._KBS50 |
| Epidermidibacterium | Epidermidibacterium_keratini |
| Marisediminicola | Marisediminicola_antarctica |
| Tessaracoccus | Tessaracoccus_defluvii |
| Changpingibacter | Changpingibacter_yushuensis |
| Yimella | Yimella_sp._cx-51 |
| Actinotignum | Actinotignum_schaalii |
| Streptosporangium | Streptosporangium_roseum |
| Dactylosporangium | Dactylosporangium_vinaceum |
| Actinocatenispora | Actinocatenispora_sera |
| Ilumatobacter | Ilumatobacter_coccineus |
| Microterricola | Microterricola_viridarii |
| Egibacter | Egibacter_rhizosphaerae |
| Streptomonospora | Streptomonospora_litoralis |
| Actinobaculum | Actinobaculum_sp._313 |
| Miniimonas | Miniimonas_sp._S16 |
| Subtercola | Subtercola_sp._PAMC28395 |
| Jiangella | Jiangella_alkaliphila |
| Nakamurella | Nakamurella_antarctica |
| Pedococcus | Pedococcus_dokdonensis |
| Actinopolymorpha | Actinopolymorpha_singaporensis |
| Friedmanniella | Friedmanniella_luteola |
| Citricoccus | Citricoccus_sp._SGAir0253 |
| Actinotalea | Actinotalea_sp._JY-7876 |
| Mumia | Mumia_sp._ZJ1417 |
| Salinibacterium | Salinibacterium_sp._ZJ450 |
| Phytohabitans | Phytohabitans_flavus |
| Haloactinobacterium | Haloactinobacterium_kanbiaonis |
| Diaminobutyricimonas | Diaminobutyricimonas_sp._LJ205 |
| Luteipulveratus | Luteipulveratus_mongoliensis |
| Kribbella | Kribbella_flavida |

| | |
|---|---|
| Stackebrandtia | Stackebrandtia_nassauensis |
| Micropruina | Micropruina_glycogenica |
| Brachybacterium | Brachybacterium_sp._CBA3105 |
| Jatrophihabitans | Jatrophihabitans_sp._GAS493 |
| Demequina | Demequina_sp._TMPB413 |
| Allobranchiibius | Allobranchiibius_sp._GilTou73 |
| Humibacter | Humibacter_sp._WJ7-1 |
| Baekduia | Baekduia_soli |
| Polymorphospora | Polymorphospora_rubra |
| Catellatospora | Catellatospora_sp._IY07-71 |
| Pseudoclavibacter | Pseudoclavibacter_sp._Marseille-Q3772 |
| Frigoribacterium | Frigoribacterium_sp._NBH87 |
| Oerskovia | Oerskovia_sp._KBS0722 |
| Herbiconiux | Herbiconiux_sp._SALV-R1 |
| Nocardia | Nocardia_farcinica |
| Gordonia | Gordonia_bronchialis |
| Tsukamurella | Tsukamurella_paurometabola |
| Tomitella | Tomitella_fengzijianii |
| Dietzia | Dietzia_psychralcaliphila |
| Skermania | Skermania_piniformis |
| Mycolicibacillus | Mycolicibacillus_parakoreensis |
| [Mycobacterium] | [Mycobacterium]_stephanolepidis |
| Hoyosella | Hoyosella_subflava |
